# Base editing of β^0^ thalassemia mutations as a therapeutic strategy for β-hemoglobinopathies: efficacy and genotoxicity studies

**DOI:** 10.1101/2025.08.21.671453

**Authors:** Giulia Hardouin, Pierre Martinucci, Samantha Scaramuzza, Panagiotis Antoniou, Federico Corradi, Alexandra Tachtsidi, Guillaume Corre, Margaux Mombled, Sandra Manceau, Cécile Rouillon, Cécile Masson, Laure Joseph, Isabelle Thuret, NathalY Network, Eric Allemand, Mario Amendola, Oriana Romano, Giuliana Ferrari, Annarita Miccio

**Author notes:** equally contributed (Mario Amendola, Oriana Romano).

## Abstract

Gene therapy has emerged as a promising curative treatment for β-hemoglobinopathies, the most common genetic disorders worldwide. However, current approved approaches still have some limitations in terms of safety and efficacy. Here, we used highly processive adenine base editors (ABE) variants to precisely correct some of the most prevalent and severe β-thalassemia-causing mutations in the β-globin gene. Efficient editing of hematopoietic stem/progenitor cells (HSPCs) led to potent β-globin expression in their erythroid progeny and persistent correction of both β-thalassemia and sickle cell-β-thalassemia phenotypes. Safety of this strategy was confirmed in HSPCs *in vitro* and *in vivo* by the absence of gene dysregulation or any meaningful impact on the DNA mutational burden, the RNA deamination level, the β-globin gene locus integrity and the clonality of the HSPC graft. Overall, base editing-mediated gene correction is a safe and effective strategy for treating β-hemoglobinopathies.

**One sentence summary:** Preclinical safety and efficacy studies of a new gene therapy approach for patients with severe β-hemoglobinopathies.

## Introduction

Beta-globin chains together with α-globin chains compose the adult hemoglobin (HbA) tetramer. β-thalassemia is a monogenic recessive disease caused by mutations in the β-globin gene (*HBB*) locus, which reduce (β^+^) or abolish (β^0^) the production of β-globin chains. The imbalance between α- and β-globin production leads to the precipitation of uncoupled α- globins, causing ineffective erythropoiesis and anemia^1^. Beta-thalassemia is a highly prevalent hemoglobinopathy with more than 42,000 affected newborns annually^2^. Patients were originally concentrated in Southeast Asia, Middle East and in the Mediterranean area but due to recent population migrations, β-thalassemia is now widely spread in Europe and North America^3^. The type of mutations influences the severity of the anemia with around 50% of patients suffering from transfusion-dependent β-thalassemia (TDT), a phenotype typically associated with β^0^/β^0^ and severe forms of β^+^ genotypes^4^.

Sickle cell β-thalassemia (BS) results from the co-inheritance of a β-thalassemic mutation and the sickle cell disease (SCD)-causing mutation. This latter mutation causes the formation of a mutant β-globin chain that causes hemoglobin (Hb) polymerization. This leads to deformation of sickle red blood cells, which occlude small vessels leading to multi-organ damage. It is the most common cause of SCD in patients of Mediterranean descent^5^. Importantly, if there is no β-globin chain production from the β-thalassemic allele (β^0^ mutation), the clinical course is similar to that of patients homozygous for the SCD mutation^5^.

While transplantation of allogenic hematopoietic stem/progenitor cells (HSPCs) represents a possible curative approach, less than 30% of patients are eligible due to the lack of compatible donors^5^. As an alternative, gene therapy approaches based on the transplantation of autologous, genetically modified HSPCs were recently approved and rely either on lentivirus (LV)- mediated gene addition or on CRISPR/Cas9-induced fetal hemoglobin (HbF) reactivation. However, while gene addition therapy has been shown to be effective for non–β^0^/β^0^ patients, β-globin transgene expression is limited and the total Hb levels tend to be lower in β^0^/β^0^ patients with iron overload and ineffective erythropoiesis often being not fully corrected^6,7^. Moreover, LV-based approaches suffer from an inherent risk of insertional mutagenesis^8^. On the other hand, CRISPR/Cas9 nuclease-based genome editing strategies^9,10^ raise safety concerns since double strand breaks (DSBs) can trigger apoptosis, genomic rearrangements and can lead to the negative selection of cells with a functional p53 pathway^11–17^.

Furthermore, while editing strategies for β-thalassemia aimed at inducing HbF have the advantage of being potentially applied to all β-thalassemia patients independently from the specific mutation, it is still debated if high HbF levels are harmful developmentally or physiologically^18^. Moreover, the FDA-approved CRISPR/Cas9 Casgevy therapy aimed at reactivating HbF leads to more variable outcomes in β-thalassemia patients who require high levels of therapeutic Hb^19^.

Therefore, development of mutation-specific gene correction approaches could allow a more predictable and physiological production of higher levels of therapeutic Hb, as the β-globin *HBB* promoter activity and *HBB* gene expression is favored in adult cells^20^.

CD39 (CAG>TAG) is one of the most common β^0^-thalassemic mutation in the Mediterranean area and Latin America, representing >40% of β-thalassemic mutations in Tunisia, Argentina and Italy^21^. It is a nonsense mutation within the codon 39, causing premature translation termination and absence of β-globin production^22^. IVS2-1 (G>A) is one of the most common β^0^-thalassemic mutations in the Middle East, representing around 30% of β-thalassemic mutations in Iran and Kuwait^23^. This point mutation disrupts the splice donor site of the second intron of *HBB* and results in the production of two abnormally spliced mRNAs, neither of which leads to the production of a functional β-globin^24^. Importantly, these severe mutations are even more represented in the group of patients requiring treatment (i.e., TDT patients)^25^.

Since the early development of genome editing tools, several approaches have been developed to correct these two prevalent β-thalassemia mutations using peptide nucleic acid (PNA) or CRISPR/Cas9 nuclease^26,27^. However, these strategies are limited by the low gene correction rate (particularly in quiescent cells, such as hematopoietic stem cells, HSCs)^26,28^ and by their toxicity (e.g., due to the delivery of a donor template carrying the wild-type β-globin sequence)^29–31^. Base editing, a new CRISPR/Cas9-derived genome editing tool, allows precise and efficient DNA modification without relying on DSB formation^32^. Adenine base editors (ABE) and cytosine base editors (CBE) contain a Cas9 nickase (nCas9) fused to a deaminase, and permit the insertion of A>G and C>T mutations, respectively^32^. Importantly, base editing has already been exploited to correct mutations causing hematopoietic disorders, demonstrating high efficiencies in *bona fide* HSCs^33–37^. We and others have previously demonstrated correction of β^+^ thalassemic mutations^35,36,38^. However, correction of the most severe β^0^ mutations and proof-of-concept of therapeutic editing and correction of the associated phenotype has not been demonstrated.

Here, we use ABEs to precisely correct the CD39 and IVS2-1 β^0^ mutations in patients HSPCs and demonstrate the rescue of the β-thalassemia and SCD phenotypes in their erythroid progeny *in vitro* and *in vivo*. Importantly, we conducted a comprehensive efficacy and safety assessment of all the potential on-target and off-target events, demonstrating the safety of ABE-based approaches for the treatment of severe β-thalassemia and sickle cell-thalassemia.

## Results

### Correction of the CD39 and IVS2-1 β^0^ mutations in β-thalassemic HSPCs results in a safe on-target editing profile

ABEs allow A•T>G•C conversions and can potentially correct the CD39 (CAG>TAG) and IVS2-1 (G>A) mutations. Therefore, we designed guide RNAs (gRNAs) placing the target base within the canonical editing window of ABEs, from positions 4 to 8 (**Supplementary Figure 1A**). For IVS2-1 mutation, we designed gRNAs that do not target a SNP close to the mutation, thus limiting the number of gRNAs to 3^39–41^ (**Supplementary Figure 1A**, rs10768683 G>C). All the gRNAs were tested in combination with the PAM-less, highly processive SpRY-ABE8e, and, when compatible, with ABE8e containing the PAM restricted nCas9 NRCH (**Table 1**).

We screened gRNA/ABE combinations by electroporating T cells obtained from β-thalassemic patients homozygous for the targeted mutations with designed gRNA/ABE combinations delivered as chemically modified gRNAs and *in vitro* transcribed ABE mRNAs (**Supplementary Figure 1B**, BT0 and BT3). gRNA1/SpRY-ABE8e and gRNA1/NRCH-ABE8e were the two most efficient combinations able to revert the CD39 mutation in more than 70% of *HBB* alleles (**Supplementary Figure 1C**). On the other hand, gRNA2/SpRY-ABE8e was the most efficient combination able to revert the IVS2-1 mutation, correcting more than 35% of the alleles (**Supplementary Figure 1C**). In all the cases, we observed low or negligible insertions/deletions (InDel) generation (**Supplementary Figure 1C**).

HSPCs from homozygous and compound heterozygous β-thalassemia patients were electroporated with selected gRNA/ABE combinations delivered as chemically modified gRNAs and *in vitro* transcribed ABE mRNAs (**Supplementary Figures 1B, Figure 1A**). For CD39, we used the NRCH-ABE8e enzyme as a more restrictive PAM requirement is expected to produce less gRNA-dependent off-targets. Deep sequencing of HSPC-derived erythroid edited samples demonstrated that gene correction was highly efficient and reproducible among the replicates (CD39: 98.1%±0.5, and IVS2-1: 90.4%±1.5), with no or few InDels (CD39: 0.0%±0.0, and IVS2-1: 0.9%±0.6) (**Figure 1B**).

**Figure 1.**
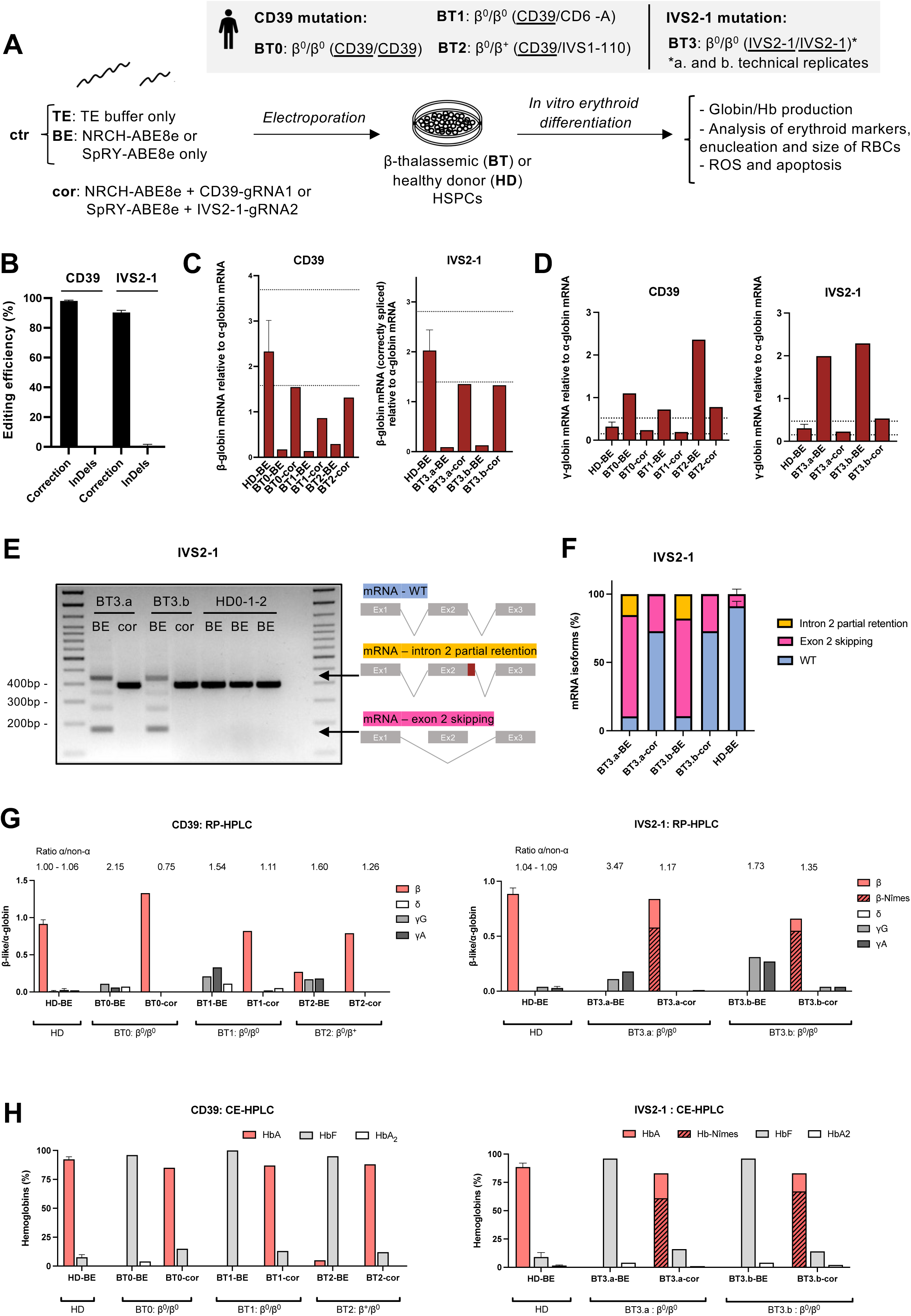
Efficient correction of the CD39 and IVS2-1 mutations in β-thalassemic HSPCs restores normal Hb production. **A.** Patients genotypes and experimental protocol used for base editing experiments in BT HSPCs. **B.** Frequency of corrected alleles and InDels as calculated after deep sequencing of *HBB* in erythroid cells derived from edited BT HSPCs. (CD39: n=3 donors; IVS2-1: 1 donor, 2 biologically independent experiments). Frequency of corrected alleles in the cells from compound heterozygous patients (BT1 and BT2) were corrected to take into account only alleles harboring the CD39 mutation. **C.** CD39: relative β-globin mRNA levels as evaluated by RT-qPCR in erythroid cells derived from corrected BT HSPCs (cor). IVS2-1: relative β-globin mRNA levels as evaluated by RT-qPCR using primers amplifying exclusively correctly spliced β-globin mRNAs in erythroid cells derived from corrected BT HSPCs (cor). As controls, we used erythroid cells derived from BT or HD HSPCs electroporated only with NRCH/SpRY-ABE8e mRNA (BE) (CD39: 3 BT, IVS2-1: 1 BT, and 3 HDs). Dotted lines indicate maximum and minimum values observed in HD cells. Of note, primers recognizing the correctly spliced β-globin mRNA hybridize to the exon2-3 junction and thus amplification of the cDNA might be impaired by the creation of the Hb-Nîmes mutation in the last codon of exon 2. Therefore, we could have underestimated the amount of correctly spliced β-globin mRNA. **D.** Relative γ-globin mRNA levels as evaluated by RT-qPCR in erythroid cells derived from control and corrected β-thalassemic HSPCs (cor). **E.** RT-PCR using primers amplifying a region from *HBB* exon 1 to exon 3. cDNA was obtained from erythroid cells derived from β-thalassemic HSPCs carrying the IVS2-1 mutation after correction (cor) (n=2 biologically independent experiments, 1 BT and 3 HDs). **F.** Relative percentages of the different isoforms of *HBB* mRNA in erythroid cells derived from β-thalassemic HSPCs carrying the IVS2-1 mutation after correction (cor). More than 6,900 reads were analyzed per sample. **E-F.** As controls, we used erythroid cells derived from BT or HD HSPCs electroporated only with SpRY-ABE8e mRNA (BE) (n=2 biologically independent experiments, 1 BT and 3 HDs). **G.** Expression of globin chains measured by RP-HPLC in BT and HD RBCs. β-like-globin expression was normalized to α-globin. The α-/non-α-globin ratio is reported on top of the graph. (CD39: n=3 donors, D13 of the differentiation; IVS2-1: n=1 donor, 2 biologically independent experiments, BT3.a= D20, BT3.b= D16 of the differentiation). **H.** Analysis of Hb variants by CE-HPLC in BT and HD RBCs. We calculated the percentage of each Hb type over the total Hb tetramers. (CD39: n=3 donors, D16 of the differentiation; IVS2-1: n=1 donor, 2 biologically independent experiments, BT3.a= D20, BT3.b= D16 of the differentiation). **G-H.** As controls, we used RBCs derived from BT or HD HSPCs electroporated only with SpRY/NRCH-ABE8e mRNA (BE). Data are expressed as mean±SEM.

Several bystander edits were observed for the CD39 gene correction strategy (**Supplementary Figure 2**, BT0-2: b0 to b4). Of note, the main bystander mutation consists in C-to-G/T mutation (**Supplementary Figure 2**, BT0-2: b3). As previously described^42^, ABE catalyzes cytosine deamination preferentially at position 6 of the protospacer within a TC*T motif (**Supplementary Figure 2**, BT0-2: b3). While correction of the mutation restores the CAG codon specifying glutamate, the simultaneous editing of the adjacent cytosine can either lead to a non-synonymous mutation (b3 = CAG>CAC: β39Gln>His) or to a synonymous mutation (b3 = CAG>CAA: β39Gln>Gln) depending on the type of base conversion (C-to-G or C-to-T, respectively; **Supplementary Figure 2**, BT0-2: b3). Importantly, β39Gln>His amino acid change (occurring in ∼4% of total alleles) has likely no clinical consequences as it was described in a known Hb variant (Hb San Bruno), which is not associated with any clinical or hematological abnormalities^43^. The four other bystander edits (all occurring in <1% of total alleles) generated non-synonymous mutations (b4 = TGG>CGG: β37Trp>Arg; b2 = AGG>AAG: β40Arg>Lys ; b1 = TTC>CTC: β41Phe>Leu or b0 = TTC>TCC: β41Phe>Ser) leading to previously described Hb variants not associated with any hematological alterations or only associated to a mild cyanosis (for β41Phe>Ser and β37Trp>Arg) (**Supplementary Figure 2**, BT0-2: b0, b1, b2 and b4 bystanders)^44–48^. In conclusion, these bystander edits occur at low frequency and are not expected impacting Hb production (see **Figure 1G-H**).

In the case of IVS2-1, two bystander mutations were observed (**Supplementary Figure 2**, BT3.a/b: b0 and b1). The first edit maps to the last codon of *HBB* exon 1 (**Supplementary Figure 2**, BT3.a/b: b0) and leads to the β104Arg>Gly missense mutation. When combined with the correction of the splicing mutation (in ∼73% of corrected alleles), this bystander edit leads to the production of a known Hb variant (Hb-Nîmes), which is not associated with any clinical abnormalities^49^. The second bystander edit (∼6% of total alleles) corresponds to an intronic known SNP (rs1468286413) with no reported clinical manifestations^50^ (**Supplementary Figure 2**, BT3.a/b: b1). However, as the IVS2-1 mutation is located within a splicing site, bystander edits could affect the splicing efficiency of *HBB* intron 2. Therefore, we used MaxEnt^51^ to predict the impact of bystander 0 and 1 on the splicing site strength. Bystander 1 (∼6% of total alleles) and to a lower extent bystander 0 (∼70% of total alleles) were predicted to decrease the strength of the splice site (**Supplementary Figure 3A**). To confirm that these bystander edits do not alter β-globin expression (and induce a β-thalassemic phenotype), we generated these base conversions in healthy donor (HD) HSPCs achieving frequencies and combinations of bystander edits similar to those detected in the BT samples (**Supplementary Figure 3A, B**). Despite a slight decrease in *HBB* expression observed at the mRNA level (**Supplementary Figure 3C**), we did not observe any effect at the protein level in HSPC-derived erythroid cells with normal HbA/Hb-Nîmes expression levels and α-/non-α globin ratio (**Supplemental Figure 3D, E**). Furthermore, differentiation was not affected, as demonstrated by the normal expression of erythroid markers (GpA, CD71 and CD36) and enucleation rate along the differentiation (**Supplemental Figure 3F, G**). Finally, generation of the bystander edits did not induce apoptosis, confirming the negligible impact of these base conversions on *HBB* expression (**Supplemental Figure 3H**).

### Efficient correction of the CD39 and IVS2-1 mutations in β-thalassemic HSPCs restores normal Hb production and corrects ineffective erythropoiesis in vitro

Following transfection, β-thalassemic HSPCs were differentiated towards the erythroid lineage (**Figure 1A**). β-globin mRNA levels in CD39 edited samples were similar to those observed in HD cells for the homozygous BT0 donor, while representing up to 50% of the HD β-globin transcripts for the compound β^0^/β^0^ heterozygote (CD39/CD6-A) and up to 80% for the compound heterozygous β^0^/β^+^ donor (CD39/IVS1-110) (**Figure 1C**). On the contrary, γ-globin mRNA expression, typically elevated in untreated thalassemic samples due to the stress erythropoiesis, was consistently reduced after treatment (**Figure 1D**). In IVS2-1 samples, RT-PCR showed restoration of the correct β-globin splicing and a substantial decrease of aberrant transcripts characterized by lower mobility due to partial intron retention or by higher mobility due to exon 2 skipping (**Figure 1E**). The relative decrease of aberrant transcripts was further confirmed by long-read sequencing of the β-globin transcripts (**Figure 1F**). The levels of correctly spliced β-globin mRNA were similar to those observed in HD cells in corrected IVS2-1 cells, while γ-globin mRNA expression was decreased in treated conditions (**Figure 1C, D**). As previously observed^52,53^, the excess of α-globin chains can be detected by RP-HPLC in thalassemic erythroid cells. Thereby, in untreated β-thalassemic RBCs, RP-HPLC showed elevated α-/non-α-globin ratios and the low β-globin expression was poorly compensated by the fetal γ (γA+γG)-globin expression (**Figure 1G**). After treatment, RBCs exhibited higher levels of β-globin chain and HbA (**Figure 1G, H)**. For IVS2-1 conditions, we detected Hb-Nîmes representing ∼64% of total Hbs (**Figure 1H**). Importantly, the α-/non-α-globin ratio was substantially ameliorated in the erythroid cells obtained from corrected β-thalassemic HSPCs (**Figure 1G**).

In β-thalassemia, α-/β-globin chain imbalance causes premature cell death via apoptosis of erythroid precursors, thus leading to ineffective erythropoiesis, a hallmark of the disease^54^. Importantly, the early erythroid markers CD36 and CD71 were properly downregulated at the end of the differentiation in samples derived from edited HSPCs and enucleation rate along the differentiation was increased, showing that the typical delayed erythroid differentiation of β-thalassemic cells was corrected by our treatment (**Figure 2A-B**). Finally, flow cytometry showed a decrease of ROS production and a rescue of edited β-thalassemic erythroid precursors from apoptosis (**Figure 2C-D**).

**Figure 2.**
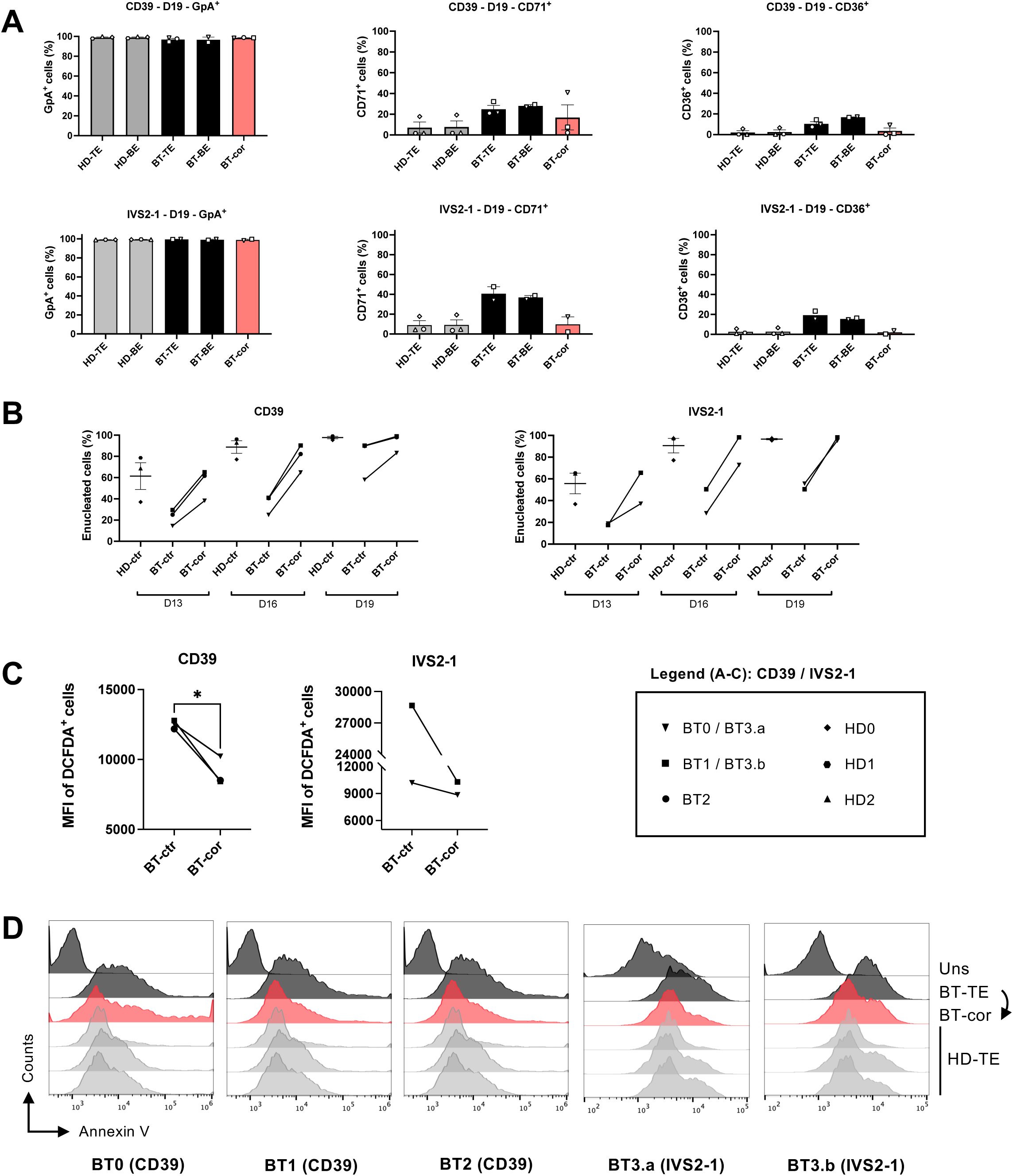
Efficient reversion of the CD39 (CAG>TAG) and IVS2-1 (G>A) mutations in β-thalassemic HSPCs corrects ineffective erythropoiesis. **A.** Frequency of GpA^+^, CD71^+^ and CD36^+^ cells at D19 of erythroid differentiation, as measured by flow cytometry analysis. As controls, we used RBCs derived from BT or HD HSPCs electroporated with TE only (TE) or with SpRY/NRCH-ABE8e mRNA only (BE) (CD39: 3 HDs and 3 BT donors; IVS2-1: n=2 biologically independent experiments, 3 HDs and 1 BT donor). **B.** Frequency of enucleated cells at D13, 16 and 19 of erythroid differentiation. **C.** Cell size of enucleated cells at D16 and D19 of erythroid differentiation, as measured by flow cytometry using the median of FSC-A intensity of DRAQ5-cells. **B-C.** As controls, we used RBCs derived from BT or HD HSPCs electroporated with TE or only with SpRY-ABE8e/NRCH-ABE8e mRNA (ctr) (CD39: 3 HDs and 3 BT donors; IVS2-1: n=2 biologically independent experiments, 3 HDs and 1 BT donor). * p≤0.05 (Paired T-test). **D.** Flow cytometry histograms showing the frequency of apoptotic cells (AnnexinV^+^-cells) in the 7AAD^-^ cell population in unstained (Uns), β-thalassemic corrected (BT-cor) or treated only with TE (BT-TE) at D13 of erythroid differentiation. As controls, we used RBCs derived from HD HSPCs electroporated with TE only (HD-TE) (CD39: 3 HDs and 3 BT donors; IVS2-1: n=2 biologically independent experiments, 3 HDs and 1 BT donor). Data are expressed as mean±SEM.

In conclusion, we were able to efficiently correct the CD39 and IVS2-1 mutations in β-thalassemic HSPCs, and to correct the pathological phenotype in HSPC-derived erythroid cells.

### Efficient correction of β-thalassemic mutations in sickle cell β-thalassemia HSPCs ameliorates the sickling phenotype in vitro

We hypothesized that targeting β-thalassemia mutations in the BS context would correct the sickling phenotype by reproducing the genotype of a SCD heterozygous carrier. Thus, we applied base editing strategies to correct the β^0^ CD39 and IVS2-1 mutations and the severe β^+^ IVS1-110 mutation^36^ in HSPCs from BS patients (**Figure 3A**). Correction of the β-thalassemia alleles was highly efficient (BS0: 90%, BS1: 83%, BS2: 81%, BS3: 91%, of total *HBB* alleles) and restored the production of high levels of therapeutic β-globin and Hb, which represented 30-50% of total Hb (**Figure 3B-C**, HbA and Hb-Nîmes). The α/non-α ratio was decreased in most of the donors upon treatment, except in BS0; we observed an increase in the α/non-α ratio after correction of the IVS2-1 β^0^ mutation and a more pronounce decrease in of HbS/βS expression compared to the donors carrying IVS1-110 or CD39 mutations, despite the similar editing efficiency (**Figure 3C**). We hypothesized that this could be (at least partially) due to aspecific targeting of the SCD allele resulting in the insertion of bystander edits responsible for a lower expression of *HBB* (**Supplemental Figure 3A**). Sequencing of *HBB* alleles revealed the insertion of the β-Nîmes mutation in SCD alleles, which could impact sickle β-globin production at mRNA and/or protein level (**Figure 3D**).

**Figure 3.**
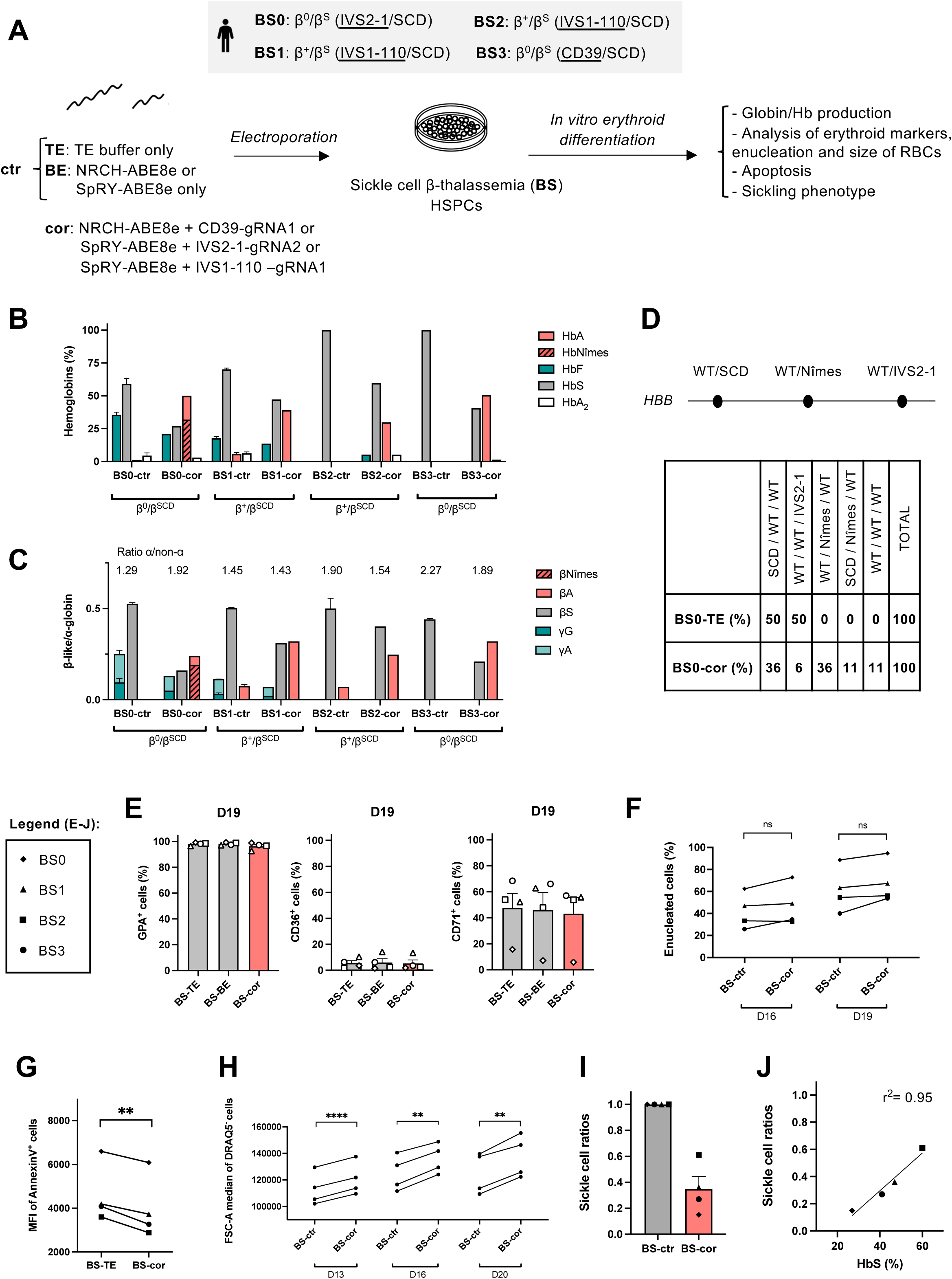
Correction of severe β-thalassemic mutations in sickle cell β-thalassemia HSPCs ameliorates the pathological phenotype *in vitro*. **A.** Experimental protocol used for base editing experiments in BS HSPCs. NRCH-ABE8e mRNA and synthetic gRNA1, SpRY-ABE8e mRNA and synthetic gRNA2 or SpRY-ABE8e mRNA and synthetic gRNA1 (previously described in^1^), were electroporated in BS HSPCs harboring the CD39, IVS2-1 or the IVS1-110 mutation, respectively. Cells were differentiated into mature RBCs using a three-phase erythroid differentiation protocol. **B.** Analysis of HbA, Hb-Nîmes, HbF, HbS and HbA_2_ by CE-HPLC in BS patients RBCs. We calculated the percentage of each Hb type over the total Hb tetramers. RBCs were obtained from corrected BS HSPCs (cor). As controls, we used RBCs derived from BS HSPCs electroporated with TE only or with SpRY/NRCH-ABE8e mRNA only (ctr). **C.** Expression of βA-, βNîmes-, βS-, γG- and γA- globin chains measured by RP-HPLC in BS patients RBCs. β-like-globin expression was normalized to α-globin. The α-/non-α-globin ratio (or mean ratio for ctr conditions) is reported on top of the graph. **D.** The region encompassing the SCD and IVS2-1 mutations in *HBB* was amplified by PCR and the PCR products were cloned and subjected to Sanger sequencing to identify the different genotypes. We reported in the table the frequency of the different *HBB* alleles retrieved in mock electroporated BS0 HSPCs (BS0-TE) and edited BS0 HSPCs (BS0-cor). **E.** Frequency of GpA^+^, CD36^+^ and CD71^+^ cells at day 19 of erythroid differentiation, as measured by flow cytometry analysis. As controls, we used RBCs derived from BS HSPCs electroporated with TE only (TE) or with SpRY/NRCH-ABE8e mRNA only (BE) (n=4 donors). **F.** Frequency of enucleated cells at day 16 and 19 of erythroid differentiation, as measured by flow cytometry analysis of cells stained with the DRAQ5 nuclear dye. **G.** MFI in apoptotic cells (Annexin^+^) in edited BS samples (cor). As controls, we used erythroid cells derived from BS HSPCs electroporated only with TE (TE). ** p≤0.01 (Paired T-test). **H.** Cell size of enucleated cells at day 13, 16 and 19 of erythroid differentiation, as measured by flow cytometry using the median of FSC-A intensity. As controls, we used RBCs derived from BS HSPCs electroporated with TE or only with SpRY-ABE8e/NRCH-ABE8e mRNA (ctr). ** p≤0.01, **** p≤0.0001 (Paired T-test). **I.** Ratio between the frequency of sickling cells (identified upon O_2_ deprivation) in RBCs derived from edited and control samples (n=4 donors). **J.** Correlation between the ratio of sickling cells and the percentage of HbS over the total Hb tetramers in RBCs obtained from corrected BS HSPCs (Simple linear regression). Data are expressed as mean±SEM.

Editing of BS HSPCs did not affect erythroid differentiation as indicated by the similar pattern of expression of erythroid markers and the comparable enucleation levels during differentiation (**Figure 3E-F**). However, correction of the β-thalassemia mutations was associated with a rescue of BS erythroid precursors from apoptosis and a larger size of enucleated erythroid cells (**Figure 3G-H**). Importantly, a deoxygenation assay (that induced Hb polymerization) showed a substantial decrease in the frequency of sickle mature RBCs obtained from base edited patients’ HSPCs compared to untreated controls (**Figure 3I**). Of note, the extent of correction of the sickling phenotype was correlated with the remaining fraction of HbS (**Figure 3J**). Overall, these results demonstrate that correction of severe β-thalassemia mutations is an efficient strategy to correct the BS phenotype.

### Correction of the β-thalassemic mutations in HSPCs: assessment of on- and off-target unwanted events

To evaluate the safety profile of our strategies, we first nominated gRNA-dependent off-target sites by *in silico* prediction^55^, and cell-based experimental cleavage detection by GUIDE-seq using the corresponding Cas9 nucleases (Cas9-NRCH and Cas9-SpRY)^56^. As GUIDE-seq performed in primary HSPCs is less sensitive in detecting off-targets than in cell lines ^57^, we electroporated K562 cells with plasmids encoding the gRNA and the Cas nucleases. We performed NGS of the top-10 *in silico* predicted (S-OTs) and top-5 empirically nominated off-target sites (E-OTs). For CD39, a total of only 10 unique off-target sites was obtained by comparing *in silico* (6 sites) and experimentally predicted sites (6 sites) (**Supplemental Figure 4A, Supplemental Table 1**). Low off-target base editing was detected at 2 sites within the β-globin gene locus (S-OT1: 9.4%±2.3 and S-OT4: 8.6%±1.0), mapping with the homologous *HBD* gene and *HBBP1* pseudogene, respectively (**Supplemental Figure 5A, Supplemental Table 1**). Importantly, the most prevalent off-target (*HBD*, S-OT1: 9.4%±2.3) has likely no consequence on the correction of the phenotype since *HBD* is poorly expressed during adulthood and the main event (CAG>CAC: *δ39*Gln>His) corresponds to a characterized SNP (rs281864505) generating a presumably neutral HbA2 variant fortuitously identified and known as HbA2-Lyon^58^. While *HBBP1* pseudogene is known to be expressed in erythroid cells and implicated in HbF regulation^59^, the off-target site does not map to any known clinically relevant SNP (i.e., associated to high HbF levels or to relevant clinical phenotypes). Lastly, we observed minimal off-target activity in an intergenic site (E-OT2: 3.3%±0.5) mapping with a region that is inactive in hematopoietic cells (**Supplemental Figure 5A, Supplemental Figure 6A, Supplemental Table 1**).

For IVS2-1, a total of 84 unique off-target sites were predicted *in silico* (42 sites) and experimentally nominated (49 sites), probably due to the lower specificity of the PAM-less ABE8e-SpRY (**Supplemental Figure 4B, Supplemental Table 1**). However, base-editing off-target events were detected exclusively in intronic (S-OT2: 90.7%±2.3, S-OT3: 8.2%±1.5, S-OT5: 2.8%±0.4), intergenic (S-OT4: 57.6%±3.9, E-OT1: 76.4%±2.6, E-OT2: 69.4%±5.0 and E-OT3: 53.9%±4.6) and lncRNA sequences (S-OT9: 6.0%±0.3) (**Supplemental Figure 5B, Supplemental Table 1**). Importantly, intronic and intergenic off-targets were located within heterochromatin or weakly transcribed regions in hematopoietic cells (**Supplemental Figure 6B**). Of note, no SNP with anticipated clinical consequence were identified in the transcribed off-target sites^60,61^. Importantly, no InDels were detected at CD39 and IVS2-1 identified off-target sites, thus minimizing the possibility of DSB-induced genomic rearrangements (e.g. translocations) (**Supplemental Figure 5A, B**). Overall, the off-target analysis combining both computational and experimental nomination methods performed for these two gene correction strategies did not identify DNA gRNA-dependent off-target mutations with anticipated clinical relevance.

Off-target editing for the CD39 gene correction strategy was detected in *HBD*, 7.5 kb away from the CD39 mutation (**Supplemental Figure 5A**, S-OT1), and this can cause genomic deletions or inversions. Therefore, we evaluated the integrity of the β-globin locus by long-read sequencing of the region encompassing the on-target site and S-OT1. As control, we transfected cells with the Cas9 nuclease and the *HBB* R-02 gRNA used in clinics in CRISPR/Cas9 nuclease-based gene correction approaches^62,63^. Long-read sequencing confirmed the integrity of the locus around the CD39 target site in the base edited patients’ sample, with no InDels (**Figure 4A-C**). On the contrary, in Cas9-treated cells, bilateral deletions were found in 70% of the sequences; large and intermediate deletions (30-200bp and >200bp respectively) accounted for a quarter of the total deletion events (**Figure 4A-C**).

**Figure 4.**
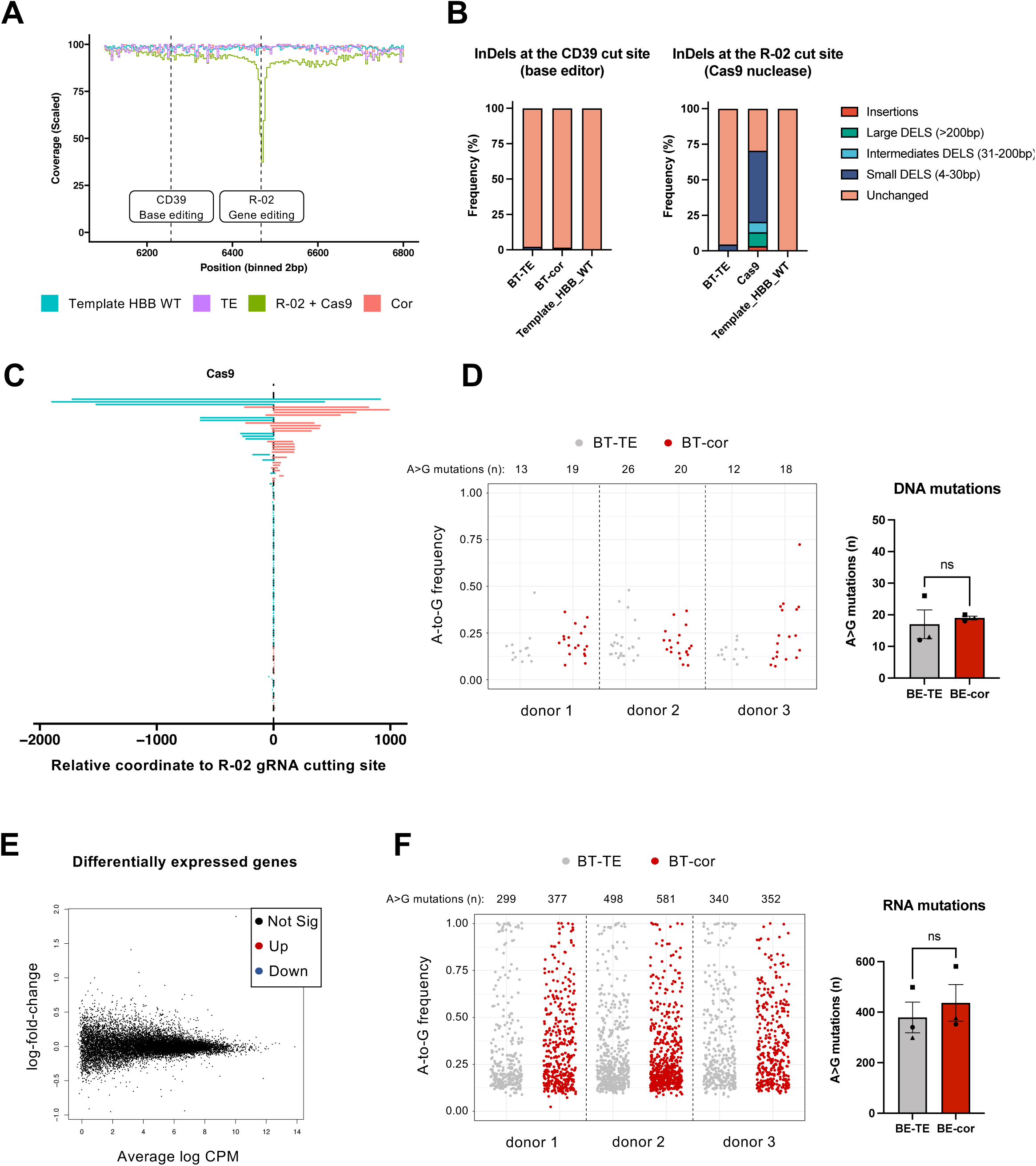
Correction of the CD39 mutation in β-thalassemic HSPCs induces no detectable genotoxic effect or transcriptomic changes. **A.** Read coverage pattern of the targeted locus in CD39 base-edited BT HSPCs (Cor). As control, we used HSPCs treated with the Cas9 nuclease in combination with R-02 *HBB* gRNA (R-02 + Cas9), mock electroporated cells (TE), as well as a library of WT reads simulated from the reference genome (Template HBB WT). **B.** Comprehensive allele frequencies of diverse InDel types induced by base editing of the CD39 mutation (left panel) or Cas9 nuclease targeting of *HBB* using the R-02 gRNA (right panel) as evaluated by long-read sequencing of the region encompassing both sites. **C.** Deletion patterns mapped relative to the cut site in HSPCs following Cas9 nuclease treatment targeting HBB R-02 site as evaluated by long-read sequencing of the locus. **D.** Strip plots showing the frequency of A>G mutations in exons as evaluated by WES of mock electroporated HSPCs (BT-TE) and corrected HSPCs (BT-cor). The total number of mutations is indicated above each sample and is plotted in the right panel. n=3 BT donors (Paired T-test, BT-TE vs BT-cor). **E.** Bulk RNA-seq analysis performed 48 h after electroporation. Mean-difference plots show differentially expressed genes between edited and mock electroporated samples. Genes that are not statistically significant (false discovery rate [FDR] > 0.05) differentially expressed are depicted with black dots. **F.** Strip plots showing the frequency of A>G mutations in RNA as evaluated by bulk RNA-seq in mock electroporated HSPCs (BT-TE) and corrected HSPCs (BT-cor). The total number of mutations is indicated above each sample and is plotted in the right panel. n=3 BT donors. (Paired T-test, BT-TE vs BT-cor).

To further evaluate the potential gRNA-independent and -dependent off-target activity in coding regions, we performed whole-exome sequencing (WES) in control and base-edited β-thalassemic HSPCs. In particular, we focused on the CD39 gene correction strategy, thanks to the availability of patients’ samples. For each donor, >99% of A>G SNVs were shared between untreated and edited samples. Regarding the remaining SNVs, we observed no significant increase in the A>G mutational burden of exons in base-edited samples compared to controls, indicating no detectable gRNA-independent off-target activity in exons (**Figure 4D**).

To uncover any unwanted effects at the RNA level, we performed whole transcriptome analysis by RNA sequencing (RNA-seq) in control and base-edited β-thalassemic HSPCs 48 h after electroporation. Importantly, we observed no differentially expressed genes showing that the base editing strategy does not affect the overall gene expression profile (**Figure 4E**). Remarkably, we did not observe dysregulation of genes belonging to the DNA damage response and the innate immune response, two pathways typically up-regulated upon Cas9 nuclease cleavage and cellular sensing of long mRNAs, respectively^17,64^. To assess the potential RNA off-target activity of our strategy, we further analyzed RNA-seq data. For each donor, >92% of A>G SNVs were shared between untreated and edited samples. The analysis of sample-specific SNVs showed that ABE8e treatment does not lead to a significant increase in the A>G endogenous RNA deamination events across the transcriptome of edited cells (**Figure 4F**).

Overall, these analyses showed that ABE8e-mediated gene correction approaches do not lead to meaningful DNA and RNA off-target activity and preserve the integrity of the on-target locus, demonstrating the safety of this strategy.

### Base editing of the CD39 mutation is maintained in repopulating HSCs and does not impair polyclonal engraftment and differentiation

To evaluate the ability of ABE8e-NRCH to correct *HBB* in repopulating HSCs, we transplanted edited mobilized human β-thalassemic HSPCs (homozygous for the CD39 mutation) into immunodeficient NBSGW mice (**Figure 5A**). HD HSPCs and unedited β-thalassemic HSPCs served as controls. Sixteen weeks post-transplantation, no significant differences were observed between edited and control HSPCs in terms of engraftment and differentiation potential, as measured by the frequency of human CD45^+^ cells and the expression of lineage-specific markers in the bone marrow (BM) (**Figures 5B-C**). Human CD45^+^ BM cells were isolated and subjected to a colony-forming cell (CFC) assay. Control and edited β-thalassemic samples showed a similar number of BFU-E, CFU-GM and CFU-GEMM, demonstrating no impact of the treatment on the clonogenic potential of engrafted human cells (**Figure 5D**). Importantly, on-target base editing was maintained at high levels over time in peripheral blood (PB) cells, and 16 weeks after transplant, in total human BM and spleen cells, in CD34^+^ HSPCs and in CD33^+^ myeloid, B lymphoid and erythroid BM subpopulations, demonstrating a successful engraftment of base edited HSCs, with no impact on HSC differentiation towards the different blood lineages (**Figure 5E**). In one mouse, we observed a lower editing efficiency in the different BM subpopulations, except in the erythroid cells (**Figure 5E**, black dot). We hypothesize that in this mouse a fraction of HSCs does not carry any corrected *HBB* allele, while corrected GpA^+^ erythroid cells possess a survival advantage over unedited cells.

**Figure 5.**
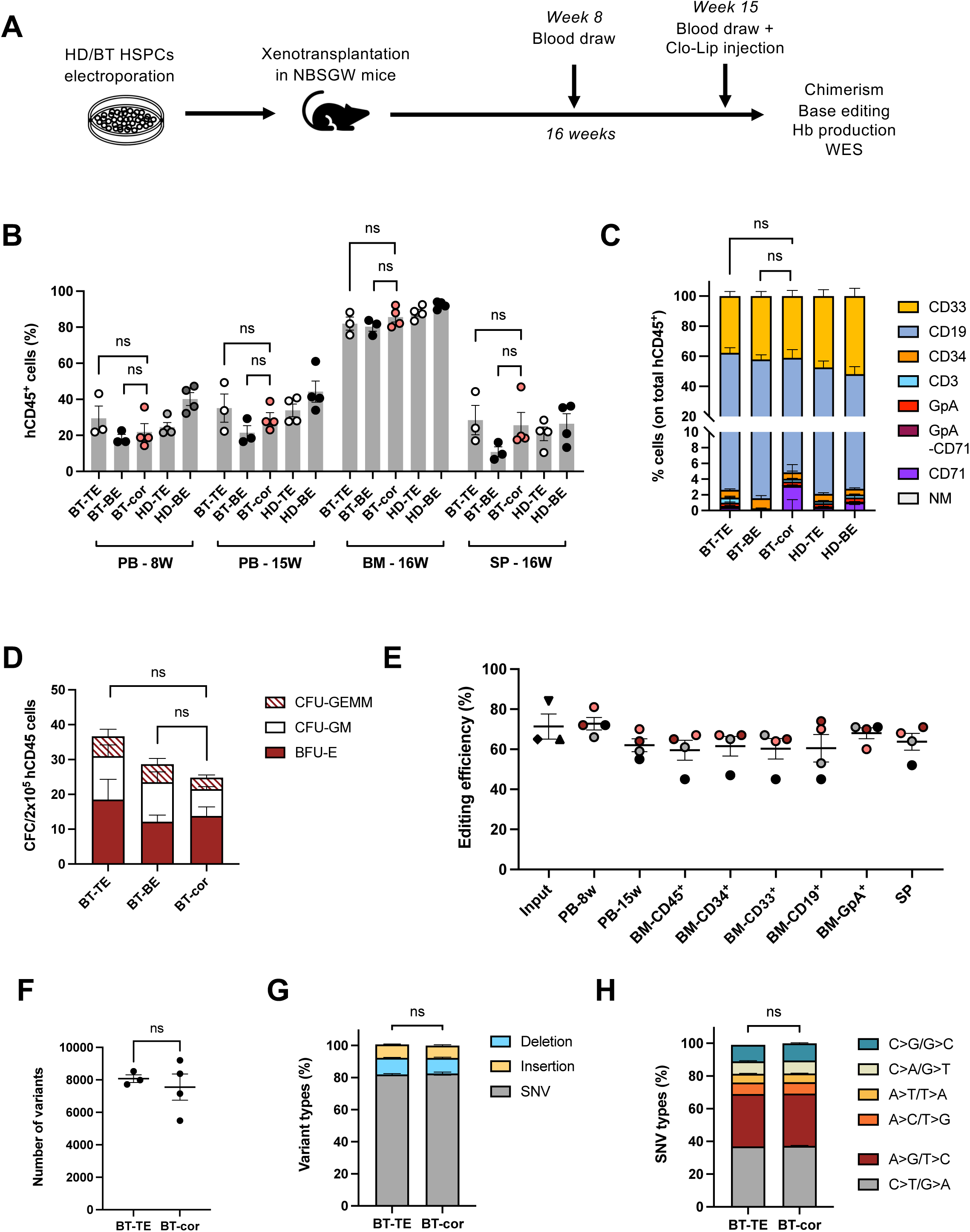
Correction of the CD39 mutation in repopulating HSCs *in vivo*. **A.** Overview of the experimental protocol of HSPC xenotransplantation. **B.** Engraftment of human cells in NBSGW mice represented as percentage of human CD45^+^ cells in the total murine and human CD45^+^ cell population in PB, BM and spleen (SP) 16 weeks post-transplantation. Mice were transplanted with BT corrected HSPCs (BT-cor) or HD/BT HSPCs electroporated with TE buffer or only with NRCH-ABE8e mRNA as controls (TE and BE conditions) (BT-TE/BE: n=3; BT-cor: n=4; HD-TE/BE: n=4). Each data point represents an individual mouse. (Unpaired T-test or Mann-Whitney test, BT-TE vs BT-cor, BT-BE vs BT-cor). **C.** Frequency of human T (CD3) and B (CD19) lymphoid, myeloid (CD33), erythroid (GpA, CD71) and HSPC (CD34) cells in BM 16 weeks post-transplantation (BT-TE/BE: n=3; BT-cor: n=4; HD-TE/BE: n=4). (Unpaired T-test or Mann-Whitney test, BT-TE vs BT-cor, BT-BE vs BT-cor). **D.** Hematopoietic progenitor content in BM human CD45^+^ cells derived from mice transplanted with control and edited HSPCs (BT-TE/BE: n=3; BT-cor: n=3). ns = non-significant (Unpaired T-test or Mann-Whitney test, BT-TE vs BT-cor, BT-BE vs BT-cor). **E.** Base editing efficiency, calculated by the EditR software, in input, PB-, BM- and SP- derived BT human samples subjected to Sanger sequencing. BT-cor: n=4. The frequency of base editing in the input was calculated in cells cultured in the HSPC medium (▾), in liquid erythroid cultures (▴), and in pooled CFU (◆) colonies. For PB, BM and SP data, each mouse is identified by a specific color. **F-H.** Number of variants (**F**), relative percentage of variants types (**G**) and relative percentage of SNV types (**H**) in the human xenograft of mice engrafted with control (TE) and corrected (cor) BT HSPCs. We subtracted the variants that were shared between untreated and treated conditions in the *in vitro* input population (TE: n=3, cor: n=4). Data are expressed as mean±SEM. Unpaired T-test or Mann-Whitney test, BT-TE vs BT-cor.

Of note, consistently with our previous observations^36^, cytosine bystander editing was higher in the CD45^+^ progeny of β-thalassemic HSCs *in vivo* compared to the HSPC input population that contains mainly hematopoietic progenitors and <1% HSCs (**Supplemental Figure 2**, BT0-BT2: b3; **Supplemental Figure 7A**). Besides C>G and C>T conversion previously described at this position in *in vitro* experiments (**Supplemental Figure 2**, BT0-BT2), we also observed a small fraction of C>A base conversion in the HSPC input population; this frequency was also substantially increased *in vivo*. The C>A conversion, however, also leads to the generation of the benign Hb San Bruno variant^43^ (b3 = CAG>CAT: β39Gln>His). We also observed a modest increase in the on-target InDel frequency in 2 out of 4 mice (**Supplemental Figure 7B**). The mouse who showed the highest InDel frequency, had also the greater frequency of C bystander edits; this observation is in line with a recent study demonstrating that InDels were more likely to occur at ABE-target sites containing TC motifs prone to C bystander editing^65^. Most of these InDels were insertions. Surprisingly, instead of being centered on the nicking site, these insertions mainly occurred around the target base, which was previously described for CBE^17^ (**Supplemental Figure 7C**).

We also evaluated editing efficiency at the two main CD39 OT sites previously described in *in vitro* experiments (**Supplemental Figure 5A,** S-OT1 and S-OT4) in total human BM cells. After 16 weeks of engraftment, we observed a decreased editing of these two sites, which was consistent with the modest reduction in the on-target gene correction efficiency. These results indicate the absence of positive or negative selection of cells carrying off-target edits (**Supplemental Figure 7B**).

We then performed WES on the input samples and the human engrafted BM cells, and used variant diversity of the clonal outgrowth as a proxy for clonality^17^. We observed a similar number of specific variants in the outgrowth of mice from the control and edited groups suggesting no impact of the base editing treatment on the clonality (**Figure 5F**). Importantly, this analysis allows us to evaluate the genome-wide effects of the editing in the progeny of long-term HSCs^17^. Interestingly, the proportion of different variant types (deletions, insertions and single-nucleotide variant; SNV) and the frequency of the different SNVs were similar between control and edited samples, demonstrating no impact of the highly processive ABE8e on the mutational landscape of edited HSCs (**Figure 5G, H**).

### Correction of the CD39 mutation in β-thalassemic HSPCs restores normal Hb production and corrects ineffective erythropoiesis in their erythroid progeny in vivo

To assess the correction of the β-thalassemic phenotype in the erythroid progeny of *bona fide* HSCs, we sorted human GpA^+^ erythroid cells from the BM of engrafted mice. Strikingly, we observed a significant increase in *HBB* mRNA expression in human erythroid cells from mice engrafted with corrected BT HSPCs (**Figure 6A**). Moreover, HPLC analysis showed the expression of β-globin in treated samples and an amelioration of the α-/non-α-globin ratio (**Figure 6B-C**). Before euthanasia, mice were subjected to clodronate liposomes (Clo-Lip) treatment in order to evaluate the egression of mature human RBCs (hRBCs) from the BM to the PB (**Figure 5A**). Interestingly, we observed a significantly increased frequency of hRBCs in the edited group of mice as compared to controls (**Figure 6D**). In the latter group, the only hRBCs that were capable to go to the PB were likely those expressing high HbF levels (as detected by HPLC), which allow them to survive and mature (**Figure 6E**). HPLC showed also the correction of the β-thalassemic phenotype in circulating hRBCs, with a high expression of β-globin and a globin expression profile overlapping with that observed in the HD control group (**Figure 6E**).

**Figure 6.**
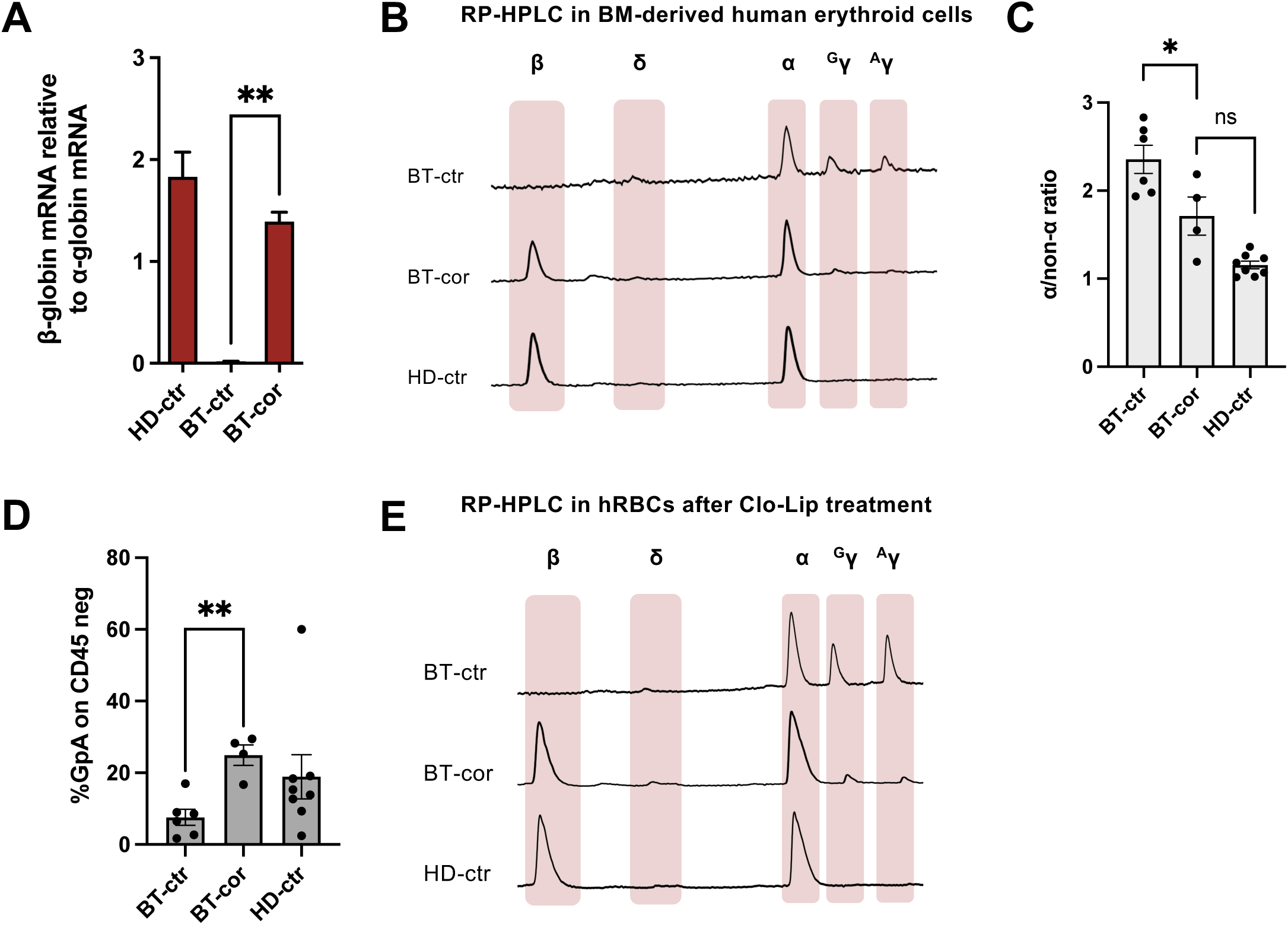
Correction of the CD39 mutation in xenotransplanted β-thalassemic HSPCs rescues the ineffective erythropoiesis *in vivo*. **A.** β-globin mRNA levels as evaluated by RT-qPCR in erythroid cells derived from the BM of mice transplanted with HD or BT control (HD-ctr; BT-ctr) or corrected (BT-cor) HSPCs (HD-ctr, n=8; BT-ctr, n=6; BT-cor, n=4). As controls, we used erythroid cells derived from β-thalassemic or HD HSPCs electroporated only with NRCH/SpRY-ABE8e mRNA (BE) (3 BT and 3 HDs). ** p≤0.01 (Mann-Whitney test, BT-ctr vs BT-cor). **B-C.** Representative RP-HPLC chromatograms of globin monomers (**B**) and α/non-α ratio (**C**) measured 16 weeks post-transplantation in sorted human GpA^+^ BM erythroid cells obtained from mice transplanted with HD or BT control (HD-ctr, BT-ctr) or corrected (BT-cor) HSPCs (BT-ctr: n=6; BT-cor: n=4; HD-ctr: n=8). * p≤0.05 (Unpaired T-test, BT-ctr vs BT-cor, BT-cor vs HD-ctr). **D.** Frequency of human RBCs in total PB 4 days after Clo-Lip injection in mice transplanted with HD or BT control (HD-ctr, BT-ctr) or corrected (BT-cor) HSPCs 16 weeks post-transplantation (BT-ctr: n=6; BT-cor: n=4; HD-ctr: n=8). ** p≤0.01 (Unpaired T-test, BT-ctr vs BT-cor). **E.** Representative RP-HPLC chromatograms from sorted human circulating RBCs 15 weeks post-transplantation.

### Clinical burden associated to mutations amenable to gene correction strategies

In order to estimate the population that could benefit from our base editing-mediated β^0^ or β^+^ mutation correction approaches (CD39, IVS1-2 and IVS1-110^36^), we queried the French National Thalassemia Registry (NaThalY) that prospectively collects laboratory and clinical data of β-thalassemia patients at national scale. As of June 2023, the database included data from 817 alive patients. Among them, 428 were TDT patients who have not received HSC transplantation at the time of study (428/817: 52.4%) (**Figure 7A**). Genotype information was available for 347 of these patients, defining our study population (347/428: 81.1%) (**Figure 7A**). Overall, a total of 181 patients carried at least one of the mutations, representing more than half of the study population (181/347: 52.2%) (**Figure 7B**). Fourteen patients were compound heterozygotes for these mutations (**Figure 7B**). CD39 was the most prevalent mutation, and was found in around 1/3 of patients (98/347), either in homozygosis (46.9%) or in combination with a β^0^ or β^+^ mutation (53.1%; **Figure 7C**). 75.5% of patients carrying the CD39 mutation were below the age of 35 years, namely the age limit of most gene therapy clinical trials (**Figure 7D**). IVS1-110 was the second most prevalent targeted mutation, carried by ∼25% of patients of the study population (86/347); one third of these patients were homozygous for the IVS1-110, while the remaining patients carry a different β^0^ or β^+^ mutation on the other allele or an α-triplication that can lead to a β-thalassemic clinical phenotype if co-inherited with a β-thalassemic mutation on only one of the two *HBB* genes (**Figure 7C**). Similarly to the CD39 population, 72.1% of patients were under 35 years (**Figure 7D**). Finally, a total of 11 TDT patients carrying the IVS2-1 mutation were identified in the study population (3.2%). The majority of them are compound heterozygous carrying this mutation in parallel with a β^0^ or β^+^ mutation (**Figure 7C**). More than 90% of these patients are under 35 years (**Figure 7D**). As the correction of only one of the two β-thalassemia alleles is sufficient to ameliorate the phenotype (**Supplementary Figure 1B, Figure 2**: BT2 and ^36^), >40% of the TDT non-HSCT French patients (146/347; carrying at least one of the targeted mutation and being under 35 years) could benefit from our base editing approaches.

**Figure 7.**
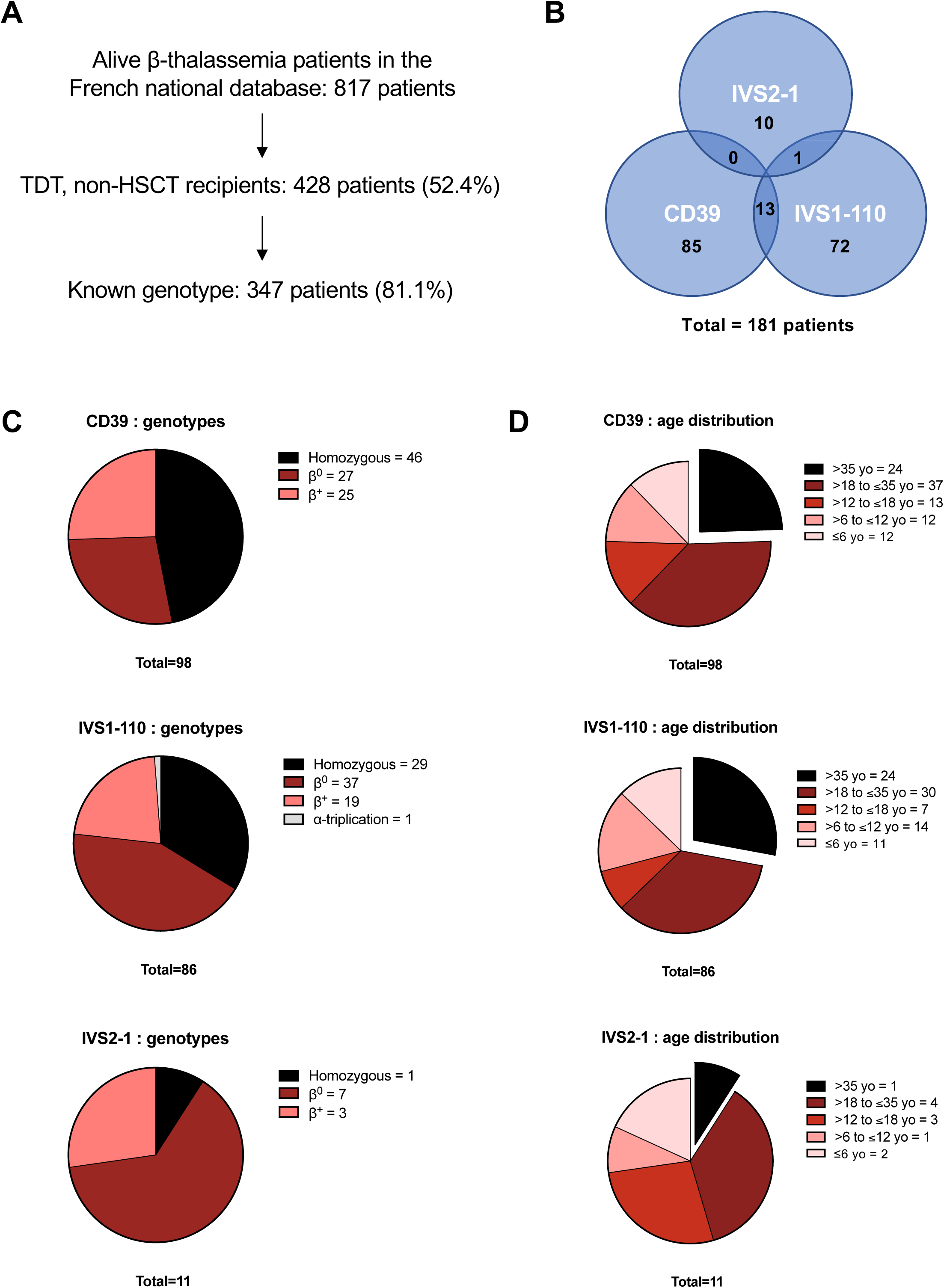
Clinical burden of mutations amenable to gene correction strategies. **A.** Flow chart describing the selection of the study population. **B.** Venn diagram representing the number of patients carrying at least one of the following mutations: CD39, IVS2-1 or IVS1-110. **C-D.** Genotype (**C**) and age (**D**) distribution in patients carrying the CD39, IVS1-110 or IVS2-1 mutation.

## Discussion

In recent years, there has been growing interest in the development of base editing strategies for the treatment of β-hemoglobinopathies^66^. Most of these approaches aim to reactivate HbF given its potential to compensate for the lack of β-globin production regardless of the causative mutation^33,64,67^. However, while the developmental and physiological impact of high HbF levels in adult life is still debated^18^, it is not yet known which strategy is the safest and most effective and future investigations comparing the safety and efficacy of both approaches should be conducted. Moreover, clinical studies with pharmacological HbF inducers suggest that treatment response could be dependent on the genetic background of individuals^68^. Similarly, the effectiveness of gene therapy approaches inducing HbF might fluctuate upon modulation by different genetic variants and could explain interindividual HbF reactivation variability^69,70^. In fact, the CRISPR/Cas9 strategies inducing HbF have shown a more variable outcome despite the similar editing efficiency^71^.

Therefore, in this study, we have developed gene correction approaches using base editors. We demonstrated that ABEs can efficiently correct two prevalent and severe β^0^ mutations in patients’ HSPCs *in vitro* and *in vivo* without relying on potentially dangerous DSBs. While nonsense-mediated decay is known to rapidly degrade CD39 β-globin mRNAs^72^, our results show the restoration of normal levels of functional mRNAs. In parallel, reversion of the β^0^ IVS2-1 mutation leads to the correction of the splicing defect and production of normal levels of correctly spliced and functional mRNAs. These strategies restored high therapeutic Hb expression in HSPC-derived RBCs *in vitro* and *in vivo*, thus correcting the β-thalassemic cell phenotype. Interestingly, we also showed evidences of *in vivo* selection of corrected human cells as previously reported in murine models^73^. Finally, to our knowledge, this study demonstrates for the first time the capacity to revert the sickling phenotype of β-thalassemia-SCD by correcting the SCD-associated β-thalassemia mutation, further expending the scope of the clinical application of our base editing strategies.

In this study, we used ABE variants derived from ABE8e, a highly-processive enzyme used in several pre-clinical studies due to its high efficiency^37,38,74,75^. Importantly, for the first time, we have provided a comprehensive and detailed efficacy and safety assessment of all the collateral on-target and off-target events. While most of the studies using base editors for therapeutic editing of HSPCs evaluated are mostly limited to gRNA-dependent DNA off target-activity, we have combined cutting-edge assays to measure efficacy and safety in terms of genotoxicity and specificity. This is particularly relevant when using the hyperactive ABE8e enzymes that, for example, could increase the frequency of unwanted on-target events^75^. Furthermore, while most of pre-clinical studies use healthy donor HSPCs as a surrogate^37,38,67^, we evaluated efficacy and safety in patients’ cells, thus taking into account the impact of the disease on the editing outcome.

First, we have characterized the effect of the different allele outcomes resulting from bystander editing. In fact, one limitation of base editing is the lack of precision due to bystander editing, which can cause unwanted events. Our study shows that, when targeting splicing junctions (IVS2-1 correction strategy), bystander edits can lead to a modest decrease expression of the target gene, which was without phenotypical consequences in cells from healthy donors or β-thalassemic patients. In contrast, the bystander editing (generating the β-Nîmes mutation) in the sickle β-globin negatively alters its expression at mRNA or protein level. By way of example this effect could be due to instability of the chimeric HbS carrying the β-Nîmes amino acid. Isolation of this chimeric Hb would be required to study its specific properties. However, the increased α/non-α ratio due to the reduced sickle β-globin production, did not result in a β-thalassemic phenotype *in vitro*. On the contrary, amongst the different β-thalassemic mutations, correction of the IVS2-1 mutation was the most effective in reducing the frequency of sickling cells, likely because of the reduction of sickle β-globin production since a lower percentage of HbS delays Hb polymerization.

Interestingly, even if the frequency was modest, base editing mediated correction of the CD39 mutation was accompanied by C>non-C bystander edits and Indels, particularly *in vivo* in long-term repopulating thalassemic HSCs (up to ∼6% of InDels in one mouse). This observation is in contrast with previous studies describing a decrease of C>non-T events (caused by CBE) and InDels following xenotransplantation of human HSPCs from healthy donors^17,33^. This suggests a potential disease-specific response to base editing in β-thalassemia HSCs. In fact, our recent study showed that β-thalassemia HSCs are less quiescent compared to those from healthy donors^77^, possibly leading to the use of different DNA repair pathways and base editing outcomes. Thus, these data highlight the need to test their consequence in the disease context in clinically relevant cells.

Importantly, it was demonstrated that cytosine deamination is not a side effect induced by adenine deamination but an independent event that could be reduced by rational design of the ABE catalytic site or by UGI addition^42,65,78^. However, modulation of the ABE activity could lead to lower A>G gene correction frequency, thus diminishing the efficiency of the strategy^42,65,78^. Bystander activity could also be reduced by using precise ABE8e variants^79,80^; however, these enzymes might have a lower on-target efficiency. Prime editing, a new CRISPR/Cas9-based technology^81^, has already been used to precisely insert CD39 and IVS2-1 mutations in erythroid cell lines and could theoretically allow the development of more precise gene correction strategies^82^, although unwanted events are also observed using this tool^83^. However, prime editing efficiency in primary hematopoietic cells is still limited^84^ and further improvements are required to use this tool for developing gene therapy approaches for hematopoietic disorders.

On-target unwanted events were also analyzed using long-read sequencing. This highlighted the interest of using base editing for gene correction strategies to preserve the β-globin gene locus integrity as compared to a CRISPR/Cas nuclease-based approach that is currently in clinics^62,63^. We showed that this latter strategy leads to unwanted deletion in the β-globin gene, while base editing mainly leads to precise gene correction.

We then comprehensively assessed the homology-dependent and homology-independent off-target activity of our base editing strategies^85^. Our study demonstrates the relevance of using Cas9 with a restrictive PAM (such as NRCH) to reduce the number of gRNA-dependent off-target sites. In fact, in the case of the ABE8e-NRCH-mediated CD39 correction strategy, the main off-target activity is restricted to low editing in two highly homologous genes of the β-globin locus. On the contrary, ABE8e-SpRY strategy is associated with a higher number of off-targets, associated with greater editing levels. Importantly, none of validated off-targets in primary HSPCs presents predictable clinical relevance, as also suggested by the absence of positive or negative selection of off-target edits *in vivo*. ABE8e with higher fidelity could be tested to reduce off-target activity, although their use is often associated with a lower on-target editing^86,87^.

WES excluded gRNA-independent off-target activity within exons in treated samples *in vitro* and *in vivo*. It has been described that CRISPR/Cas9 nuclease treatment leads to a reduced clonality *in vivo*^17,88^. Here in our work, WES of the clonal outgrowth of human cells engrafted in immunodeficient mice showed no impact of ABE8e treatment on the clonality and mutational burden of the graft. We believe that this is particularly important given the high mutational burden that can be observed in patients with β-hemoglobinopathies and that can be associated to a high incidence of clonal hematopoiesis^89^. Finally, despite the use of the hyperactive ABE8e variants, RNA-seq experiments in patients HSPCs showed no increase in RNA deamination in edited samples (i.e., gRNA-independent RNA off-target activity) compared to untreated controls. Furthermore, RNA-seq experiments showed that base editing has a negligible impact on the transcriptome of patients HSPCs.

To our knowledge, this extended genotoxicity analysis demonstrates for the first time the safety and efficacy of the ABE8e highly processive enzyme usage in HSCs. Our results support the further development of base editing approaches for HSCs based on this enzyme.

While publicly available data highlight the clinical burden of the selected mutations in the Mediterranean area^21,23,90^, we extracted precise demographic and medical data from the French National Thalassemia Registry and observed that around 50% of TDT non-HSCT patients carry one of the mutations amenable to ABE (CD39, IVS1-110 and IVS2-1). When considering current age limitations for a gene therapy procedure, we estimated that >40% of TDT patients who have not received an allogeneic HSCT could benefit from such a treatment. Of note, while being represented in our study center, the IVS2-1 mutation is not broadly present in France. Additional studies combining demographic and clinically relevant information in other regions where these mutations are even more prevalent could be of interest to better evaluate the population that could benefit from these approaches. For instance, a similar study conducted in Italy showed that more than 80% of TDT patients carry CD39 or IVS1-110 mutation^25^. Altogether, these gene correction approaches are of particular interest given the high percentage of β-thalassemia patients who could benefit and their clinical development could pave the way for similar approaches for other mutations that are prevalent in other geographic areas.

Overall, our study demonstrates that gene correction using base editing is safe and efficient in *bona fide* patient’s HSCs, thus paving the way for the clinical development of these approaches for the treatment of β-thalassemia and sickle-cell-thalassemia.

## Material and Methods

### HSPC and T cell purification and culture

We obtained human non-mobilized PB CD34^+^ HSPCs from β-thalassemia and sickle cell β-thalassemia patients (for in vitro experiments). Samples eligible for research purposes were obtained from the “Hôpital Necker-Enfants malades” Hospital (Paris, France) or “Hôpital de la Timone” (Marseille, France). Written informed consent was obtained from all adult subjects. All experiments were performed in accordance with the Declaration of Helsinki. The study was approved by the regional investigational review board (reference: DC 2022-5364, CPP Ile-de-France II “Hôpital Necker-Enfants malades”). HSPCs were purified by immunomagnetic selection immunostaining with the CD34 MicroBead Kit (Miltenyi Biotec). The CD34^-^ fraction was kept for T cell cultures.

Plerixafor/GCSF-mobilized PB CD34+ cells (used for in vivo experiments) were selected from a patient affected by TDT β-thalassemia upon signed informed consent approved by the Ethical Committee of the San Raffaele Hospital (Milan, Italy). Following mobilization and apheresis, CD34+ cells were purified using immunomagnetic beads (CliniMACS, Miltenyi Biotec) by MolMed SpA (Milan, Italy)^91^ and cryopreserved.

Forty-eight hours before electroporation, CD34^+^ cells were thawed and cultured at a concentration of 5×10^5^ cells/mL in the “HSPC medium” containing StemSpan (STEMCELL Technologies) supplemented with penicillin/streptomycin (Gibco), 250 nM StemRegenin1 (STEMCELL Technologies), and the following recombinant human cytokines (PeproTech): human stem cell factor (SCF) (300 ng/ml), Flt-3L (300 ng/ml), thrombopoietin (TPO) (100 ng/ml), and interleukin-3 (IL-3) (60 ng/ml). Four days before electroporation, the CD34^-^ fraction was thawed and cultured at 5×10^6^ cells/mL in the “T cells medium” containing RPMI 1640 + GlutaMAX (Gibco) supplemented with FBS (Thermo), penicillin/streptomycin (Gibco) and Recombinant Human IL-2 (Peprotech). After recovery, cells were transferred to “T cell activation medium” supplemented with CD28 Monoclonal Antibody (eBioscience, Clone CD28.2) in plates coated with CD3 Monoclonal Antibody (eBioscience, Clone OKT3).

### Base editor plasmids

Constructs used in this study include NRCH-ABE8e and SpRY-ABE8e plasmids. The NRCH-ABE8e plasmid was created by replacing the Cas9-coding sequence of the ABE8e plasmid (Plasmid #138489, Addgene)^92^ with the Cas9-NRCH included in the “pCMV-ABEmax-NRCH” plasmid (Plasmid #136923, Addgene)^93^. The SpRY-ABE8e plasmid was created by replacing the Cas9-coding sequence of the ABE8e plasmid (Plasmid #138489, Addgene)^92^ with the Cas9 fused to GFP included in the “pCMV-T7-ABEmax(7.10)-SpRY-P2A-EGFP (RTW5025)” plasmid (Plasmid #140003, Addgene)^94^.

### gRNA design

We manually designed gRNAs targeting the CD39 and IVS2-1 mutations (**Supplemental Table 2**). The gRNA used to target the IVS1-110 (G>A) mutation was previously described^36^. We used chemically modified synthetic gRNAs harboring 2′-O-methyl analogs and 3′-phosphorothioate nonhydrolyzable linkages at the first three 5′ and 3′ nucleotides (Synthego).

### mRNA in vitro transcription

20 μg of NRCH-ABE8e or SpRY-ABE8e expressing plasmids were digested overnight with SapI restriction enzyme (Thermo) that cleaves once right after the poly-A tail. The linearized plasmids were purified using a PCR purification kit (QIAGEN #28106) and were eluted in 14 μl of DNase/RNase-free water. 2 μg of linearized plasmid were used as template for the *in vitro* transcription reaction (MEGAscript, Ambion #AM1334). The *in vitro* transcription protocol was modified as follows. The GTP nucleotide solution was used at a final concentration of 3.0 mM instead of 7.5 mM and the anti-reverse cap analog N7-Methyl-3’-O-Methyl-Guanosine-5’-Triphosphate-5’-Guanosine (ARCA, Trilink #N-7003) was used at a final concentration of 12.0 mM resulting in a final ratio of Cap:GTP of 4:1 that allows efficient mRNA capping. The incubation time for the *in vitro* reaction was reduced to 30 minutes. mRNA was precipitated using lithium chloride and resuspended in TE buffer in a final volume that allowed to achieve a concentration of >1 μg/μl. The mRNA quality was assessed using Bioanalyzer (Agilent).

### RNA electroporation

1×10^6^ T cells per condition were electroporated with 3.0 μg of the ABE-encoding mRNA and3.2 μg of the synthetic gRNA. When ABE was not fused to GFP, a GFP-encoding mRNA (Tebu-bio) was added to the transfection mix. We used the P3 Primary Cell 4D-Nucleofector X Kit S (Lonza) and the EO115 program (Nucleofector 4D). Cells electroporated only with TE buffer served as negative controls.

For *in vitro* experiments, 1×10^4^ to 5×10^5^ HSPCs per condition were electroporated with 3.0 μg of the ABE-encoding mRNA and 3.2 μg of the synthetic gRNA. We used the P3 Primary Cell 4D-Nucleofector X Kit S (Lonza) and the CA137 program (Nucleofector 4D). Cells electroporated only with TE buffer or only with the ABE-encoding mRNA served as negative controls.

For *in vivo* experiments, 1.25 to 2.5 ×10^6^ HSPCs per condition were electroporated with 15.0 μg of the ABE-encoding mRNA and 16.0 μg of the synthetic gRNA using the P3 Primary Cell 4D-Nucleofector X Kit L (Lonza) and the CA-137 program (Nucleofector 4D). Cells electroporated only with TE buffer or only with ABE-encoding mRNA served as negative controls.

### HSPC differentiation

Electroporated CD34^+^ HSPCs were differentiated into mature red blood cells (RBCs) using a three-phase erythroid differentiation protocol, as previously described^95,96^. During the first phase (day 0 to day 6), cells were cultured in a basal erythroid medium supplemented with 100 ng/ml recombinant human SCF (PeproTech), 5 ng/ml recombinant human IL-3 (PeproTech), 3 IU/ml EPO Eprex (Janssen-Cilag) and 10^−6^ M hydrocortisone (Sigma). During the second phase (day 6 to day 9), cells were co-cultured with MS-5 stromal cells in the basal erythroid medium supplemented with 3 IU/ml EPO Eprex (Janssen-Cilag). During the third phase (day 9 to day 22), cells were co-cultured with stromal MS-5 cells in a basal erythroid medium without cytokines. Erythroid differentiation was monitored by flow cytometry analysis of CD36, CD71, GYPA and of enucleated cells using the DRAQ5 double-stranded DNA dye. 7AAD was used to identify live cells.

### Evaluation of editing efficiency

Base editing efficiency and InDel frequency were evaluated in HSPC-derived erythroid cells at D6 or D13 of differentiation, and in T cells 3 days after electroporation. Genomic DNA was extracted from control and edited cells using PURE LINK Genomic DNA Mini kit (LifeTechnologies), or Quick-DNA/RNA Miniprep (ZYMO Research), following manufacturers’ instructions. To evaluate base editing efficiency in T cells, we performed either a nested PCR using previously published primers for CD39^97^, or a regular PCR with newly designed primers for IVS2-1 (Fw: GGATCTGTCCACTCCTGATGC, Rv: ACTGTACCCTGTTACTTATCCCC) followed by Sanger sequencing and EditR analysis^98^. TIDE analysis (Tracking of InDels by Decomposition) was performed to evaluate the percentage of InDels in edited samples^99^.

On-target and off-target regions of DNA from HSPC-derived erythroid cells or hCD45^+^ sorted mice BM cells were PCR-amplified and subjected to NGS. Off-targets were nominated using COSMID^55^ and by GUIDE-seq^56^. We selected the top-10 *in silico* predicted and top-5 GUIDE-seq predicted off-target sites and assessed editing at day 9 or 13 of differentiation. On-target and off-target sites were PCR-amplified using the Phusion High-Fidelity polymerase (NEB, M0530) and primers containing specific DNA stretches (MR3 for forward primers and MR4 for reverse primers; **Supplemental Table 3**) located 5’ to the sequence recognizing the off-target. For the on-target site of CD39, a nested PCR was performed. Amplicons were purified using Ampure XP beads (Beckman Coulter, A63881). Illumina-compatible barcoded DNA amplicon libraries were prepared by a second PCR step using the Phusion High-Fidelity polymerase (NEB, M0530) and primers containing Unique Dual Index (UDI) barcodes and annealing to MR3 and MR4 sequences. Libraries were pooled, purified using the High Pure PCR Product Purification Kit (Sigma-Aldrich, 11732676001), and sequenced using Illumina NovaSeq 6000 system (paired-end sequencing; 2×100-bp) to obtain a minimum of 15,000 reads per amplicon. Targeted NGS data were analyzed using CRISPResso2^100^.

For evaluating the frequency of cell harboring both SCD and Nimes mutations, the *HBB* allele was amplified with the following primers encompassing the SCD mutation and IVS2-1 mutation loci: FW= TCCCCTTCCTATGACATGAACTT, RV= TGCTTACATTTGCTTCTGACACA. PCR products were cloned into the pCR-XL-2-TOPO vector following the manufacturer instructions (K8050-10, Invitrogen). A minimum of 34 colonies were sequenced per condition.

### In silico splicing prediction

Wild-type, mutant and edited *HBB* gene sequences were analyzed using the MaxEntScan::score5ss for human 50 splice sites^51^ based on the maximum entropy principle (http://hollywood.mit.edu/burgelab/ maxent/Xmaxentscan_scoreseq.html).

### GUIDE-seq

DNA library preparation for GUIDE-seq analysis was performed as previously described^56,101^. Human lymphoblast K562 cells (2.5×10^5^) were transfected with 4 μg of Cas9-coding, 800 ng of gRNA-coding plasmid and 80 pmol of double-stranded oligonucleotide containing phosphorothioate bonds at both ends. After 5 days in culture, DNA was isolated with QIAamp DNA mini kit using standard protocols (QIAGEN). DNA fragments of 400–900 bp were generated by sonication using a Bioruptor Pico sonicator and subsequently ligated to adaptors, followed by two steps of DNA amplification by utilizing KAPA HyperPrep Kit (Roche KK8504). Then, the libraries were purified and measured using the Qubit fluorometer (Thermo Fisher Scientific), the concentrations were evaluated using the qPCR KAPA libraries quantification kit (Roche) and the average bp length was estimated using TapeStation Bioanalyzer 2100 (Agilent). Finally, the libraries were pooled, diluted to reach a concentration of 4 nM and loaded into a MiSeq flow cell. Demultiplexing, PCR duplicate consolidation, cleavage on-target site recognition, off-target activity identification, and visualization was performed with the GUIDE-Seq Analysis pipeline from Bushman lab (https://github.com/cnobles/iGUIDE) using the hg38 human genome as reference and RefSeq for annotations.

### Long-read sequencing of HBB by Cas9-enrichment library preparation

gRNAs were designed to introduce cuts on complementary strands flanking the region of interest (ROI), using CHOPCHOP online design tool (https://chopchop.cbu.uib.no/) and selected for the highest predicted on-target efficiency and minimal off-target activity. The gRNAs were assembled as a duplex from synthetic CRISPR RNAs (crRNAs) (Integrated DNA Technologies-IDT, custom designed) and trans-activating crRNAs (tracrRNAs) (IDT). gRNA sequences are provided in **Supplemental Table 4**. The ROI was centered on the expected cut sites of the gene editing approaches on *HBB* gene and has a size of ∼14 kb. High molecule weight DNA was extracted from HSPC patient cells with the Nanobind CBB kit (PacBio) according to manufacturer’s instructions. DNA was size selected using Short Read Eliminator XS kit (PacBio) and quantified using the Qubit fluorometer (Thermo Fisher Scientific). 5 µg of DNA were used for the library preparation using the Cas9 Sequencing Kit (Oxford Nanopore Technologies-ONT, SQK-CS9109) and following the Cas9-mediated PCR-free enrichment protocol (version: CAS_9106_v109_revC_16Sep2020) available through ONT. Libraries were loaded onto PromethION flow cells with R9.4.1 nanopores (ONT). One flow cell was used per biological condition and run on a P2 solo using MinKNOW software for 72h. Raw reads from Nanopore sequencing were preprocessed using cutadapt (v4.4) to remove low quality bases (q=5 threshold at 5’ and 3’ ends) and select reads longer that 4 kb. Adaptors were then trimmed using porechop (v 0.2.4) and reads aligned on the human reference genome (GRCh38.p13) using minimap2 (v 2.26-r1175; arguments*: -x map-ont -a -Y --secondary=no*).

Reads that align at the enrichment locus (11:5220519-55121185) were re-aligned on this region using minimap2 (arguments*: -x map-ont -n 20 -I 1K -r 400,1000 -k 15 -w 15 -a -Y -- secondary=no).* Finally, the depth of coverage at each position was measured using samtools (v1.17), the base composition using pysamstats (v1.1.2) and the presence of InDels detected using the variant caller sniffles (v 1.0.12; arguments*: -n -1 -r 1000 -s 1 -d 1 -l 4*). Additional processing was performed using R (v4.0.2) to prepare figures (ggplot2 & ggseqlogo libraries) and tables. InDels were classified as small, intermediate and large according to their size ([4-30bp], [31-200bp], >200bp). Fastq files containing Nanopore sequencing data have been deposited in the ENA database (https://www.ebi.ac.uk/ena/browser) under accession code PRJEB78636.

### Whole exome sequencing

For *in vitro* experiments, genomic DNA was extracted 6 days following electroporation (Purelink Genomic DNA mini kit, Invitrogen). Exome libraries were prepared by Imagine Genomic Core Facility (Paris, France) using Twist Human Core Exome (36.8 Mb, Twist Bioscience). Briefly, genomic DNA was sheared with an Ultrasonicator (Covaris). A total amount of 30 ng of the fragmented and purified double strand DNA was used to prepare Twist Exome libraries as recommended by the manufacturer. Barcoded exome libraries were pooled and sequenced with the Illumina NovaSeq X system (paired-end sequencing; 2×150-bp) by the NGS service provider Macrogen. More than 75 million paired-end reads were produced per exome library.

For *in vivo* experiments, genomic DNA was extracted from input populations 6 days following electroporation, or from the human cells isolated from xenotransplanted mice at the time of euthanization (QIAamp Mini/Micro Kit, Qiagen). Exome libraries were prepared using Twist Human Core Exome + RefSeq (Twist Bioscience) and sequenced with the Illumina NovaSeq X system (paired-end sequencing; 2×150-bp) by the NGS service provider Macrogen. More than 96 million paired-end reads were produced per exome library.

Read quality was evaluated using FastQC (v. 0.11.9)^102^. Raw reads were trimmed for adapters and low-quality tails (quality < Q20) with BBDuk (v. 38.92)^103^. Reads shorter than 35 bp after trimming were removed.

Variant calling was carried out accordingly to GATK Best Practices for germline short variant discovery (GATK v4.2.2.0)^104^. In brief, FASTQ files were mapped on the hg38 human reference genome with BWA (v 0.7.17)^105^, specifying the ReadGroup. For samples retrieved from mice (in vivo dataset), to remove possible mouse contaminations an in silico Combined human-mouse Reference Genome (ICRG; hg38 + mm10)^106^ was used to discriminate between human and mouse reads; then only reads assigned to the hg38 human reference genome were retained. Duplicates were marked using GATK MarkDuplicates. Base quality recalibration was performed using GATK BaseRecalibrator and ApplyBQSR, specifying the list of target exons with a padding region of 100 bp. Variant calling was performed using GATK HaplotypeCaller only on canonical (1–22, X, Y and M) chromosomes. SNVs and InDels were hard-filtered using GATK VariantFiltration applying suggested basic thresholds from GATK Best Practices. SNVs annotation was performed using the Variant Effect Predictor (VEP) tool from Ensembl^107^. Multiallelic variants (mainly involving repetitive sequences) were removed. To identify private variants belonging to each sample, only SNVs with coverage ≥ 20 reads and genotype quality ≥ 30 were retained. For the in *vitro* dataset, to define SNVs private to the treated sample a frequency of the reference allele ≥ 0.99 was required in the untreated sample, and vice versa. For the *in vivo* dataset, to define variants present in the original cell population prior to any manipulation (germline variants), shared variants between untreated and treated input samples were retrieved and removed from all the bone marrow derived samples. Fastq files containing WES have been deposited in the SRA database (https://www.ncbi.nlm.nih.gov/sra) under accession code PRJNA1144227 (in vitro dataset: https://dataview.ncbi.nlm.nih.gov/object/PRJNA1144227?reviewer=2r4dhd1lrc75kqs62ae5c2 itla) and PRJNA1144228 (in vivo dataset: https://dataview.ncbi.nlm.nih.gov/object/PRJNA1144228?reviewer=1o5hgbv58tajjjk4avhp9d ac3o).

### RT-PCR and RT-qPCR

For *in vitro* experiments, RNA was extracted from cells at day 13 of differentiation (Qiagen, 74004 or Zymo Research, ZD7001) and retro-transcribed (Thermo, 18080051). For *in vivo* experiments, total RNA was extracted (ALLPREP DNA/RNA Micro Kiyt, Qiagen) from GpA^+^ cells purified from BM and PB of transplanted immunodeficient mice and retrotranscribed with SuperScript IV VILO cDNA Synthesis Kit with EzDNase treatment (Thermo Fisher) following manufacturer’s instructions. RT-PCR analysis of globin mRNAs was performed using previously described primers spanning introns 1 and 2^108^. RT-qPCR was performed using previously described primers for the detection of correctly spliced β-globin mRNA^97^ and the following primers: α-globin-F 5’-CGGTCAACTTCAAGCTCCTAA-3’, α-globin-R 5’-ACAGAAGCCAGGAACTTGTC-3’, β-globin-F 5’-GCAAGGTGAACGTGGATGAAGT-3’, β-globin-R 5’-TAACAGCATCAGGAGTGGACAGA-3’, γ-globin-F 5’-CCTGTCCTCTGCCTCTGCC-3’, γ-globin-R 5’-GGATTGCCAAAACGGTCAC-3’.

### Long-read sequencing of HBB mRNA transcripts

Total RNAs (600 ng) were subjected to reverse transcription (RT) with an anchored poly-dT primer and Maxima H Minus Reverse Transcriptase using the following reaction conditions: 2 minutes at 37°C, 30 minutes at 50°C, 20 minutes at 65°C, and 5 minutes at 85°C. The amplification of the *HBB* transcriptome was independently processed in each sample through two PCR steps, using 2 μL of diluted RT reactions (12-fold dilution). First, a pre-amplification was performed for 20 cycles using specific primers fused with universal sequences U1 and U2 (U1-HBB: U1-GACACAACTGTGTTCACTAGCAACCTC; U2-HBB: U2-GGACAGCAAGAAAGCGAGCTTAGTG). These primers are specifically base paired with the first and last exons of *HBB* transcripts. The PCR reactions were treated with exonuclease I to remove primer excess and purified with the NucleoMag® NGS Clean-up and Size Select (Macherey-Nagel). Next, a second amplification of 18 cycles was performed to incorporate barcodes associated with individual samples. All samples were combined to create a stoichiometric multiplexed library, which was prepared using the Oxford Nanopore SQK-LSK109 kit. The library was subsequently sequenced using a MinION Flow Cell (R9.4.1). The Oxford Nanopore data were analyzed as previously described^109^ and Fastq files containing Nanopore sequencing data have been deposited in the ENA database (https://www.ebi.ac.uk/ena/browser) under accession code PRJEB78471.

### RNA-seq

Total RNA was isolated from edited HSPCs 48 h following electroporation (ZD7001, Zymo Research). RNA quality and concentrations were assessed by capillary electrophoresis using High Sensitivity RNA reagents with the Fragment Analyzer (Agilent Technologies).

Bulk mRNA-seq libraries were prepared by the Imagine Genomic Core Facility (Paris, France) starting from 50 ng of total RNA using the NEBNext® Single Cell/Low Input RNA Library Prep Kit according to manufacturer’s guidelines. This kit generates mRNA-seq libraries from the PolyA+ fraction of the total RNA by template switch reverse transcription. The cDNAs were amplified twice and the final mRNA-seq libraries were ‘unstranded’. The equimolar pool of libraries (assessed by Q-PCR KAPA Library Quantification kit, Roche and with a run test using the iSeq100, Illumina) was sequenced on Illumina NovaSeq6000 (S4 FlowCell, paired-end sequencing, 2x150-bp). More than 60 millions of paired-end reads were produced per library.

Read quality was evaluated using FastQC (v. 0.11.9)^102^. Raw reads were trimmed for adapters and low-quality tails (quality < Q20) with BBDuk (v. 38.92)^103^. Moreover, the first 10 nucleotides were force-trimmed for low quality. Reads shorter than 35 bp after trimming were removed. Trimmed reads were aligned to the human reference genome (hg38) using STAR (v. 2.7.9a)^110^. Raw gene counts were obtained in R-4.1.1 using the featureCounts function of the Rsubread R package (v. 2.8.1)^111^ and the GENCODE v44 basic gene annotation for hg38 human reference genome. Gene counts were normalized to counts per million mapped reads (CPM) and to fragments per kilobase of exon per million mapped reads (FPKM) using the edgeR R package (v. 3.36.0)^112^; only genes with a CPM greater than 1 in at least 3 samples were retained. Differential gene expression analysis was performed using the glmQLFTest function of the edgeR R package, using donor as a blocking variable.

RNA editing analysis was performed accordingly to GATK Best Practices for RNA-seq variant calling (GATK v4.2.2.0)^104^. In brief, FASTQ files were two-pass aligned to the hg38 human reference genome with STAR (v. 2.7.9a)^110^ using parameters to specify the ReadGroup and output the aligned BAM file sorted by coordinate; then duplicates were marked using GATK MarkDuplicates. After splitting reads containing Ns in their cigar string because they span splicing sites using GATK SplitNCigarReads, base quality recalibration was performed using GATK BaseRecalibrator and ApplyBQSR, and the known variants collected in dbSNP155. RNA base-editing variant calling was performed using GATK HaplotypeCaller only on canonical (1–22, X, Y and M) chromosomes. Single-nucleotide variants (SNVs) were hard-filtered using GATK VariantFiltration applying suggested basic thresholds from GATK Best Practices. SNVs annotation was performed using the Variant Effect Predictor (VEP) tool from Ensembl. Multiallelic variants (mainly involving repetitive sequences) were removed. To identify private variants belonging to each sample, only SNVs with coverage ≥ 20 reads and genotype quality ≥ 30 were retained. Moreover, to define SNVs private to the treated sample a frequency of the reference allele ≥ 0.99 was required in the untreated sample, and vice versa. Fastq files and gene expression matrix containing RNA-seq data were deposited in the GEO database (https://www.ncbi.nlm.nih.gov/geo/query/acc.cgi?acc=GSE273814; accession code GSE273814, enter token qncromuypzsdjix into the box).

### Flow cytometry analysis

Flow cytometry analysis of CD36, CD71 and GYPA erythroid surface markers on HSPC-derived erythroid cells was performed using a V450-conjugated anti-CD36 antibody (561535, BD Horizon), a FITC-conjugated anti-CD71 antibody (555536, BD Pharmingen) and a PE-Cy7-conjugated anti-GYPA antibody (563666, BD Pharmingen). Flow cytometry analysis of enucleated or viable cells was performed using double-stranded DNA dyes (DRAQ5, 65-0880- 96, Invitrogen and 7AAD, 559925, BD, respectively). Apoptosis was evaluated using PE Annexin V Apoptosis Detection Kit I (BD Biosciences). Flow cytometry analysis of reactive oxygen species (ROS) was performed using H_2_DCFDA (D3999, Invitrogen). Flow cytometry analyses were performed using Gallios (Beckman coulter) flow cytometer. Data were analyzed using the FlowJo (BD Biosciences) software.

### RP-HPLC analysis of globin chains

Reversed-phase HPLC analysis was performed using a NexeraX2 SIL-30AC chromatograph and the LC Solution software (Shimadzu). A 250x4.6 mm, 3.6 μm Aeris Widepore column (Phenomenex) was used to separate globin chains by HPLC. Samples were eluted with a gradient mixture of solution A (water/acetonitrile/trifluoroacetic acid, 95:5:0.1) and solution B (water/acetonitrile/trifluoroacetic acid, 5:95:0.1). The absorbance was measured at 220 nm.

### CE-HPLC analysis of hemoglobin tetramers

Cation-exchange HPLC analysis was performed using a NexeraX2 SIL-30AC chromatograph and the LC Solution software (Shimadzu). A 2 cation-exchange column (PolyCAT A, PolyLC, Columbia, MD) was used to separate hemoglobin tetramers by HPLC. Samples were eluted with a gradient mixture of solution A (20mM bis Tris, 2mM KCN, pH=6.5) and solution B (20mM bis Tris, 2mM KCN, 250mM NaCl, pH=6.8). The absorbance was measured at 415 nm.

### Sickling assay

HSPC-derived mature RBCs obtained at the end of the erythroid differentiation, were incubated under gradual hypoxic conditions (20% O2 for 20min; 10% O2 for 20min; 5% O2 for 20min; 0% O2 for 120-150 min) and a time course analysis of sickling was performed in real-time by video microscopy. Images were captured using an AxioObserver Z1 microscope (Zeiss) and a 40x objective and then processed with ImageJ to determine the percentage of sickle RBCs per field of acquisition in the total RBC population. More than 200 cells were counted per condition.

### HSPC xenotransplantation in NBSGW mice

NOD.Cg-KitW-41JTyr +PrkdcscidIl2rgtm1Wjl/ThomJ (NBSGW) were bred and maintained in a specific pathogen-free (SPF) animal facility. Procedures were performed according to protocols approved by the Committee for Animal Care and Use of San Raffaele Scientific Institute (Committee Protocol # 1203). 6-9 weeks old mice were conditioned with a non-lethal dose of busulfan intraperitoneally (15 mg/Kg body weight) and 24 hours after, mice were injected with 0.2 x 10^6^ G-CSF/Plerixafor-mobilized PB cells from control or β-thalassemic cells via retro-orbital sinus injection. 15 weeks after transplantation, mice were injected intraperitoneally with 10 ug/kg clodronate liposomes (Liposoma). 4 days after, 30 uL of PB were collected and analyzed using the following antibodies: anti-human CD45-VioBlue, (Miltenyi Biotec), anti-human CD235/GpA-FITC (Biolegend), anti-human CD19-PECy7 (BD), 1/50 anti-human CD3-BV510 (Biolegend), anti-human CD71-APC (BD Pharmingen). GpA expressing cells were purified by immunomagnetic selection with CD235/GpA Microbeads (Miltenyi Biotec). 16 weeks after HSPC infusion, mice were euthanized, and organs (BM and spleen) were collected. Cells from PB and organs were harvested and analyzed for the presence of human CD45^+^ (using an anti-human CD45-VioBlue antibody, Miltenyi Biotec) cells and the expression of different lineage specific surface markers (using an anti-human CD235/GpA FITC antibody (Biolegend); an anti-human CD33-APC antibody (Miltenyi Biotec); an anti-human CD19-PECy7 antibody (BD); an anti-human CD3-BV510 (Biolegend); an anti-human CD34-PE antibody (BD); an anti-human CD71 APC antibody (BD Pharmingen); and analyzed by CANTO II flow-cytometer (BD) and FCS Express v7 Software (De Novo Software). Human CD45^+^ cells from BM were sorted by immunomagnetic negative selection with murine cells depletion Kit (Miltenyi Biotec). Human CD19^+^, CD34^+^, CD33^+^ and GpA^+^ cells were sorted from BM cells by FACS Aria Fusion Cell Sorter (BD).

### CFC assay

CFC assay was performed using the Methocult medium (H4434, STEMCELL Technologies) from *in vitro* treated CD34^+^ cells plated at the concentration of 500 and 1000 cell/ml and collected 14 days after for evaluation of base editing efficiency. CFC assay was also performed from human cells from BM of transplanted mice (after depletion of murine cells) plated at the concentration of 0.15 x 10^6^ cells/mL and counted 14 days after.

## List of Supplementary Materials

Fig. S1 to S7

Table S1 to S4

## Acknowledgments

The authors would like to thank Christine Bole and the Imagine genomic facility for the generation of the next-generation sequencing (NGS) data and WES library preparation. We are grateful to the patients for their participation.

## Author contributions

G. H. designed, conducted the experiments, analyzed data and wrote the paper. P. M., S. S., P. A., F. C., A. T., M. M., C. R. and E. A. conducted the experiments and analyzed data. G. C. and E.A analyzed long-read sequencing data. G. C. analyzed GUIDE-seq data. C. M. analyzed off-target data. O. R. analyzed WES and RNA-seq data. M. A. contributed to the design of the experimental strategy. S. M., L. J. and G. F. contributed to the design of the experimental strategy and provided patients’ samples. I. T. and the NathalY Network provided epidemiological data and patients’ samples. A. M. conceived the study and wrote the paper.

## Data and materials availability

All data needed to evaluate the conclusions in the paper are present in the paper and/or the Supplementary Materials. Fastq files containing WES have been deposited in the SRA database under accession code PRJNA1144227 and PRJNA1144228. Fastq files and gene expression matrix containing RNA-seq data were deposited in the GEO database; accession code GSE273814, enter token qncromuypzsdjix into the box). Fastq files containing Nanopore sequencing data have been deposited in the ENA database under accession codes PRJEB78471 and PRJEB78636. Additional data related to this paper may be requested from the authors.

## Supplemental figure legends

**Supplementary Figure 1.**
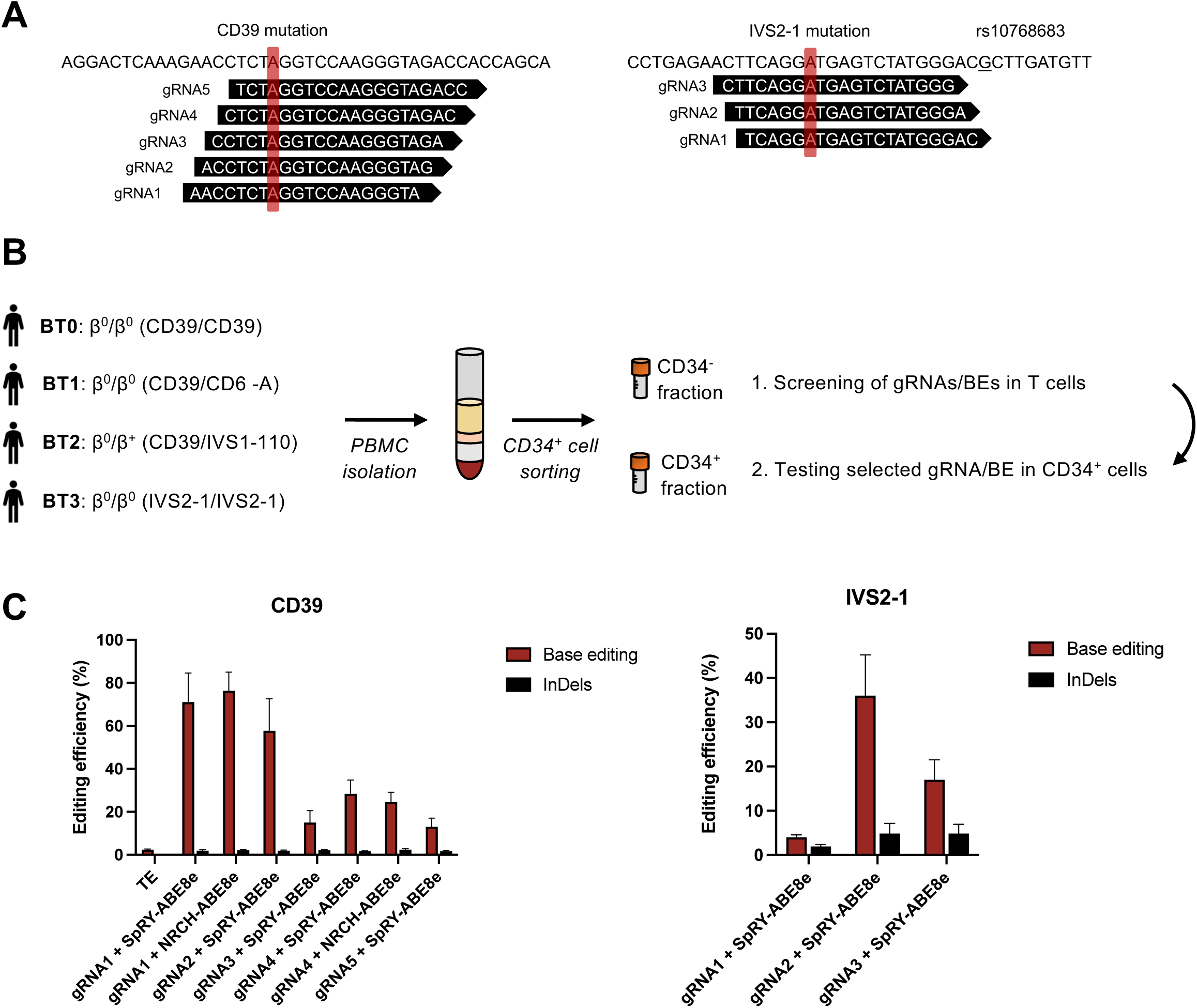
Design and screening of gRNAs targeting the CD39 and IVS2-1 mutations in β-thalassemic T cells. **A.** CD39 and IVS2-1 gRNAs sequences. The mutation is highlighted with a red box. The rs10768683 SNP proximal to the IVS2-1 mutation is underlined. **B.** Overview of the cell collection process. PBMCs were isolated from one homozygous (BT0) and two compound heterozygous (BT1 and BT2) β-thalassemia patients harboring the CD39 mutation and 1 homozygous patient harboring the IVS2-1 mutation (BT3, BT3.a and BT3.b when used in duplicates). **C.** Frequency of corrected alleles as evaluated by EditR and InDel frequency as assessed by TIDE in T cells electroporated with different combinations of synthetic gRNAs and ABE mRNAs. n=3 biologically independent experiments, 1 homozygous donor per mutation. Data are expressed as mean±SEM.

**Supplementary Figure 2.**
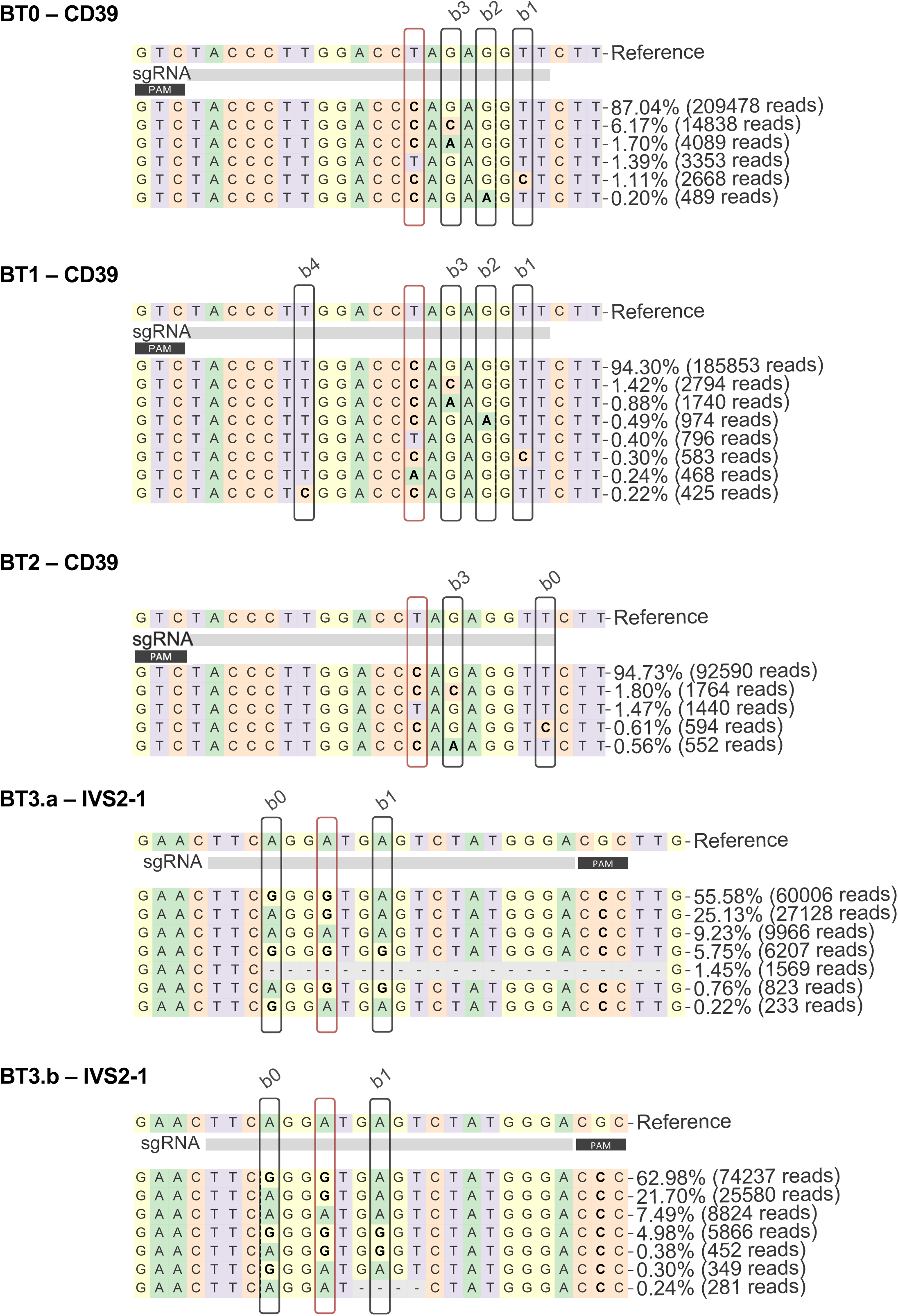
On-target editing in β-thalassemic HSPCs. Frequency and sequence of modified and unmodified alleles in erythroid cells derived from β-thalassemic HSPCs, as measured by targeted NGS sequencing. The target base position is highlighted with a red box. Bystander edits are highlighted with black boxes.

**Supplementary Figure 3.**
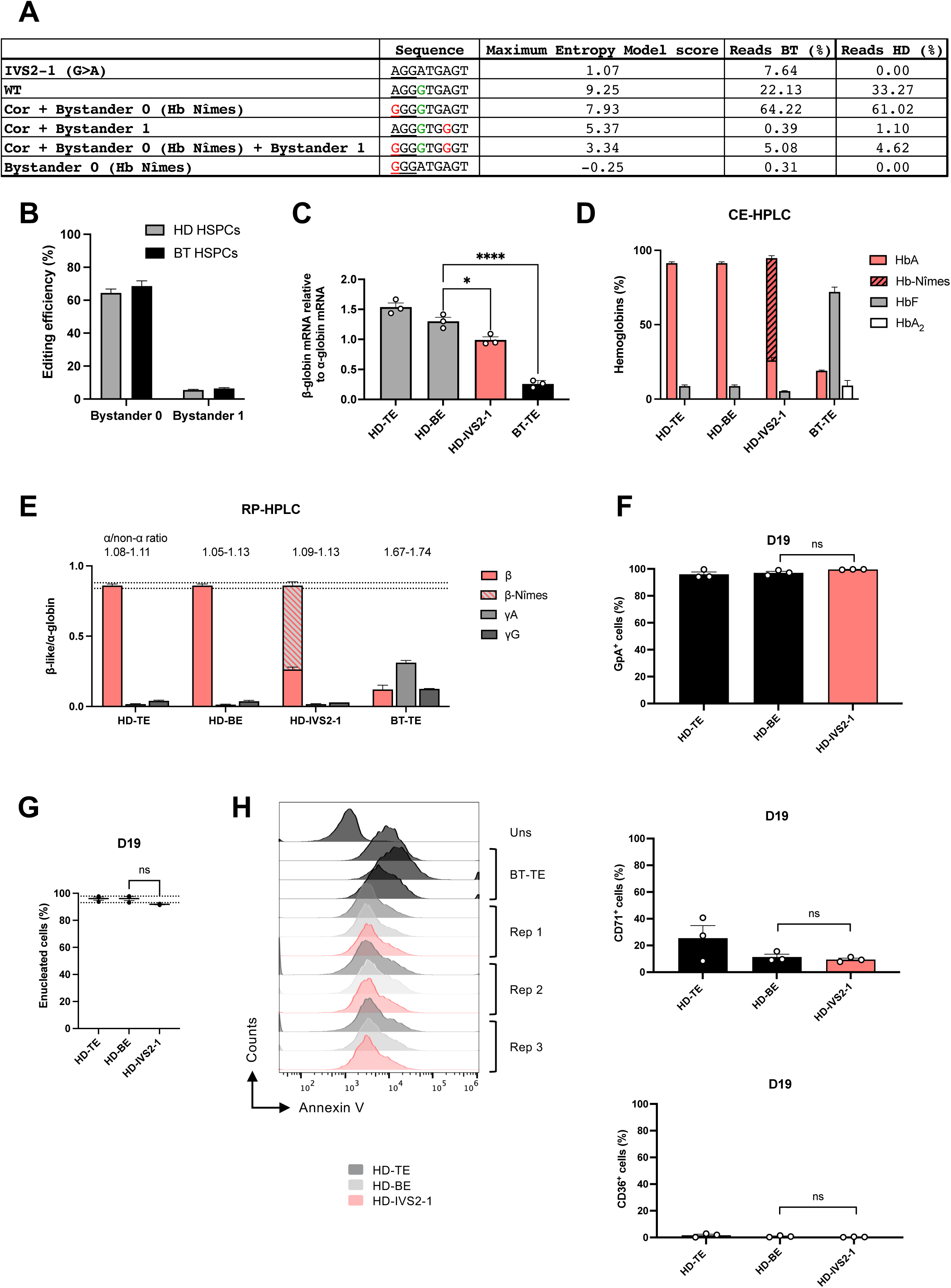
Impact of bystander edits on *HBB* expression. **A.** Sequences, mean percentage of reads in BT and HD HSPCs and scores of the 5’ splice site as predicted using the tool MaxEntScan::score5ss for human 5’ splice sites, with the Maximum Entropy Model. The higher is the score, the stronger is the 5’ splice site sequence. The corrected base is depicted in green; bystander edits are depicted in red. **B.** Frequency of alleles carrying bystander 0 and bystander 1 in edited HD and BT samples (n=3). Data are expressed as mean±SEM. **C.** RT-qPCR detecting total β-globin mRNAs in erythroid cells derived from edited HD cells carrying the bystander edits (HD-IVS2-1). As controls, we used erythroid cells derived from HD HSPCs electroporated only with TE (HD-TE) or SpRY-ABE8e mRNA (HD-BE). Data from erythroid cells derived from control BT HSPCs were included (BT-TE; 3 BT patients). β-globin expression was normalized to α-globin. Data are expressed as mean±SEM (n=3). * p≤0.05, **** p≤0.0001 (One-way Anova, HD-BE vs other conditions). **D.** Analysis of HbA, Hb-Nîmes, HbF and HbA2 by CE-HPLC in HD RBCs. We calculated the percentage of each Hb type over the total Hb tetramers. RBCs were obtained from edited HD cells carrying the bystander edits (HD-IVS2-1). As controls, we used erythroid cells derived from HD HSPCs electroporated only with TE (HD-TE) or SpRY-ABE8e mRNA (HD-BE). Data from erythroid cells derived from BT HSPCs were included (BT-TE; 3 BT patients). Data are expressed as mean±SEM (n=3). **E.** Expression of β-, β-Nîmes-, ^A^γ- and ^G^γ- globin chains measured by RP-HPLC in HD RBCs. β-like-globin expression was normalized to α-globin. The α-/non-α globin ratio is reported on top of the graph. RBCs were obtained from edited HD cells carrying the bystander edits (HD-IVS2-1). As controls, we used erythroid cells derived from HD HSPCs electroporated only with TE (HD-TE) or SpRY-ABE8e mRNA (HD-BE). Data from erythroid cells derived from BT HSPCs were included (BT-TE; 3 β-thalassemia patients). Data are expressed as mean±SEM (n=3). Dotted lines indicate maximum and minimum values observed for β-globin in HD-TE/BE conditions. **F.** Frequency of GpA^+^, CD71^+^ and CD36^+^ cells at day 19 of erythroid differentiation, as measured by flow cytometry analysis. RBCs were obtained from edited HD cells carrying the bystander edits (HD-IVS2-1). As controls, we used erythroid cells derived from HD HSPCs electroporated only with TE (HD-TE) or SpRY-ABE8e mRNA (HD-BE). Data are expressed as mean±SEM (n=3). Paired t-test (HD-BE vs HD-IVS2-1). **G.** Frequency of enucleated cells at day 19 of erythroid differentiation, as measured by flow cytometry analysis of cells stained with the DRAQ5 nuclear dye. RBCs were obtained from edited HD cells carrying the bystander edits (HD-IVS2-1). As controls, we used erythroid cells derived from HD HSPCs electroporated only with TE (HD-TE) or SpRY-ABE8e mRNA (HD-BE). Data are expressed as mean±SEM (n=3). Dotted lines indicate maximum and minimum values of enucleation observed in HD-TE/BE conditions. Paired t-test (HD-BE vs HD-IVS2-1). **H.** Flow cytometry histograms showing the frequency of apoptotic cells (AnnexinV^+^-cells) in the 7AAD-cell population of HD samples carrying the bystander edits (HD-IVS2-1) at day 13 of erythroid differentiation. As controls, we used erythroid cells derived from HD HSPCs electroporated only with TE (HD-TE) or SpRY-ABE8e mRNA (HD-BE). Data from erythroid cells derived from BT HSPCs were included (BT-TE). Uns = unstained, rep = replicate (n=3).

**Supplementary Figure 4.**
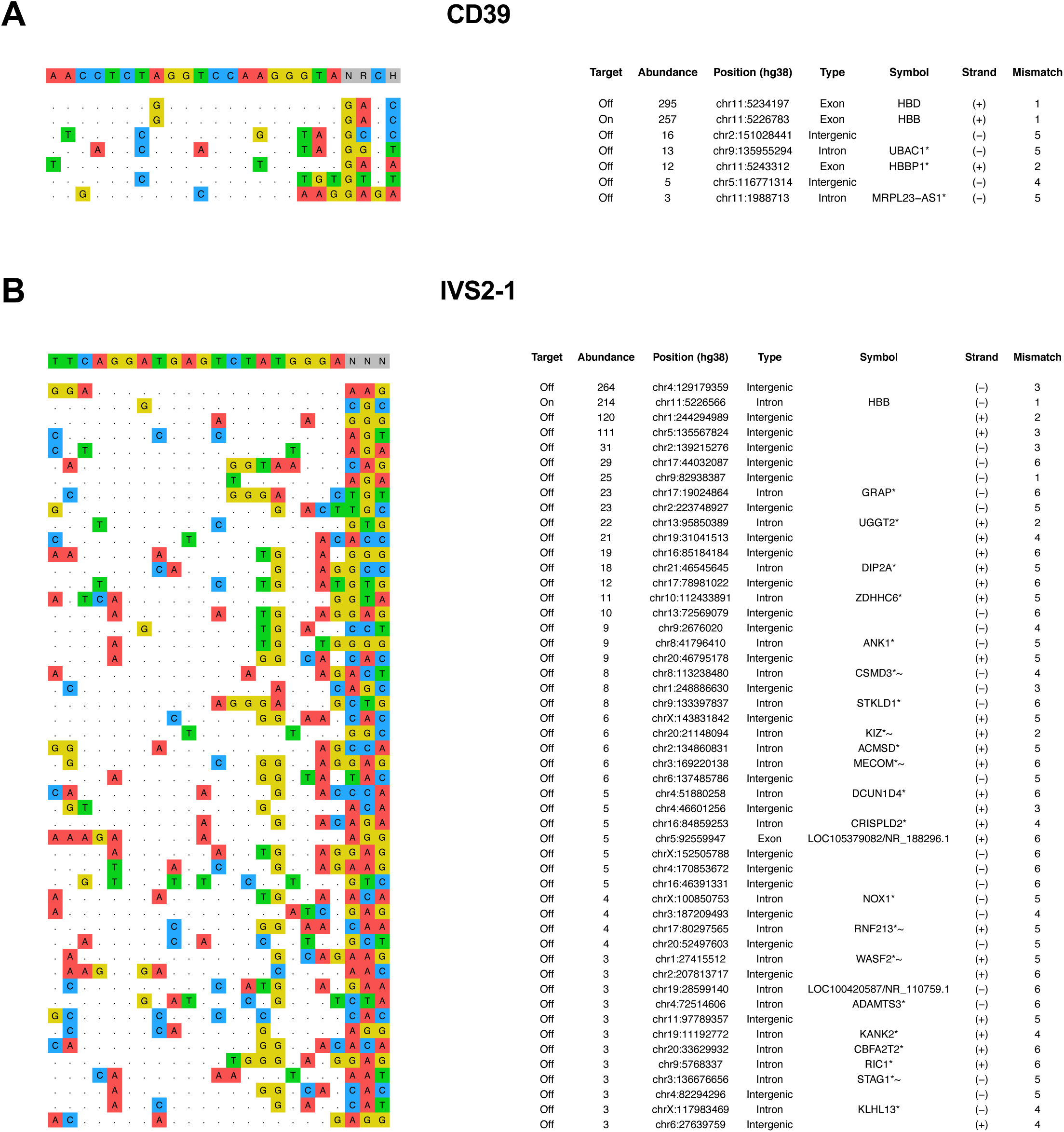
GUIDE-seq analysis of gRNA-dependent DNA off-target activity. **A-B.** gRNA-dependent off-target sites of the CD39 gRNA1 (**A**) and the IVS2-1 gRNA2 (**B**) in K562 cells, as evaluated by GUIDE-seq analysis. gRNAs were coupled with a Cas9-nuclease corresponding to the Cas9 nickase included in the base editor (Cas9-NRCH for CD39 gRNA1 and Cas9-SpRY for IVS2-1 gRNA). The protospacer targeted by each gRNA and the PAM are reported on top of each panel, followed by the off-target sites and their mismatches with the on-target (highlighted in color). The number of sequencing reads, the chromosomal coordinates (hg38), and the site of each off-target are reported. Cutoff was set to a level of detection of at least 3 reads. (*) indicates that the site is within the transcription unit of the gene, and (∼) indicates that the gene appears in the cancer-association list.

**Supplementary Figure 5.**
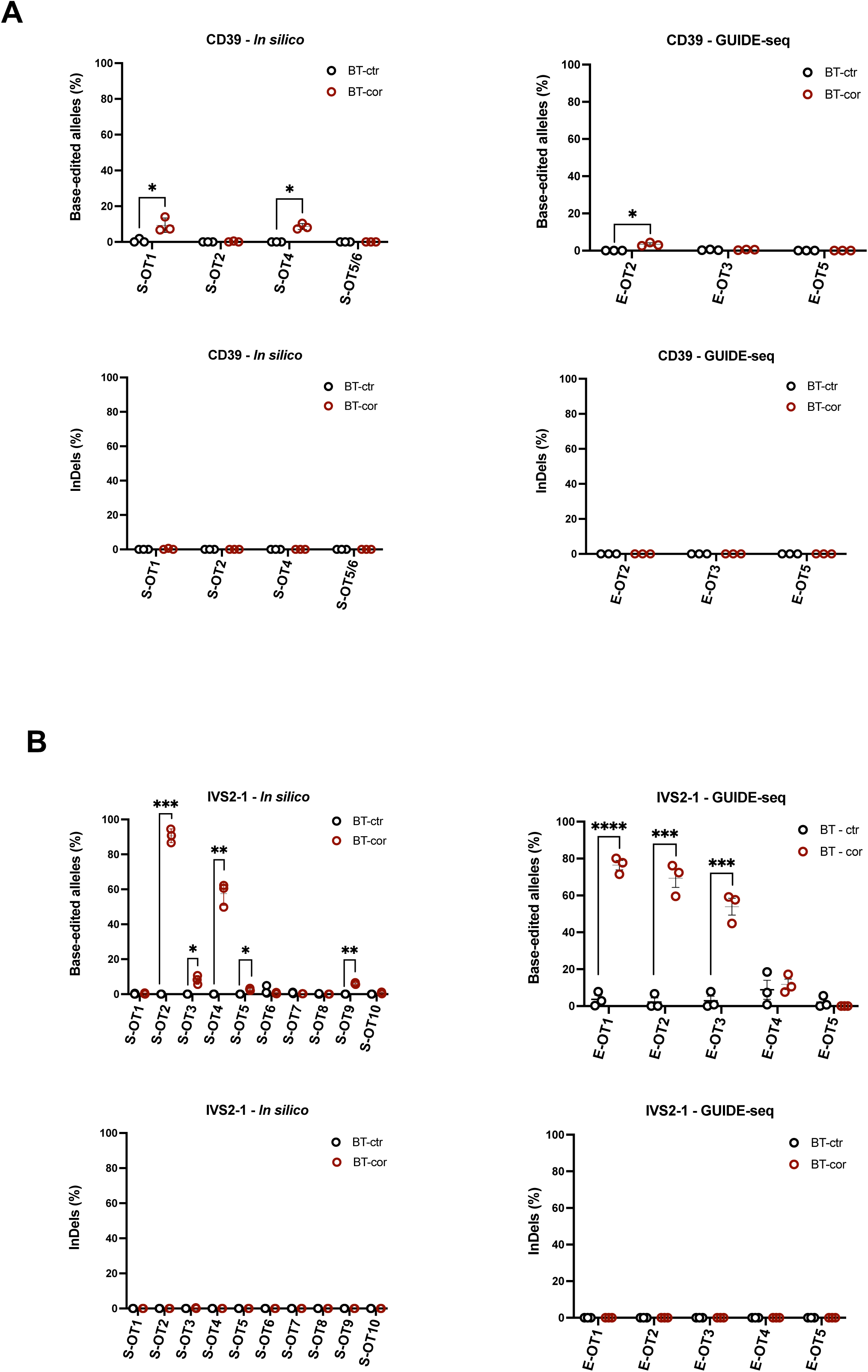
Off-target editing in β-thalassemic cells. Frequency of base editing and InDels at the top-10 predicted (*in silico*) and top-5 empirically nominated (GUIDE-seq) off-target sites in control (BT-ctr) and edited (BT-cor) β-thalassemic samples, as measured by targeted NGS sequencing in BT-HSPCs following editing of the CD39 (**A**) or IVS2-1 (**B**) mutation (n=3 donors). * p≤0.05, ** p≤0.01, *** p≤0.001, **** p≤0.0001 (Unpaired t-test). Of note, CD39 S-OT3 site could not be PCR-amplified.

**Supplementary Figure 6.**
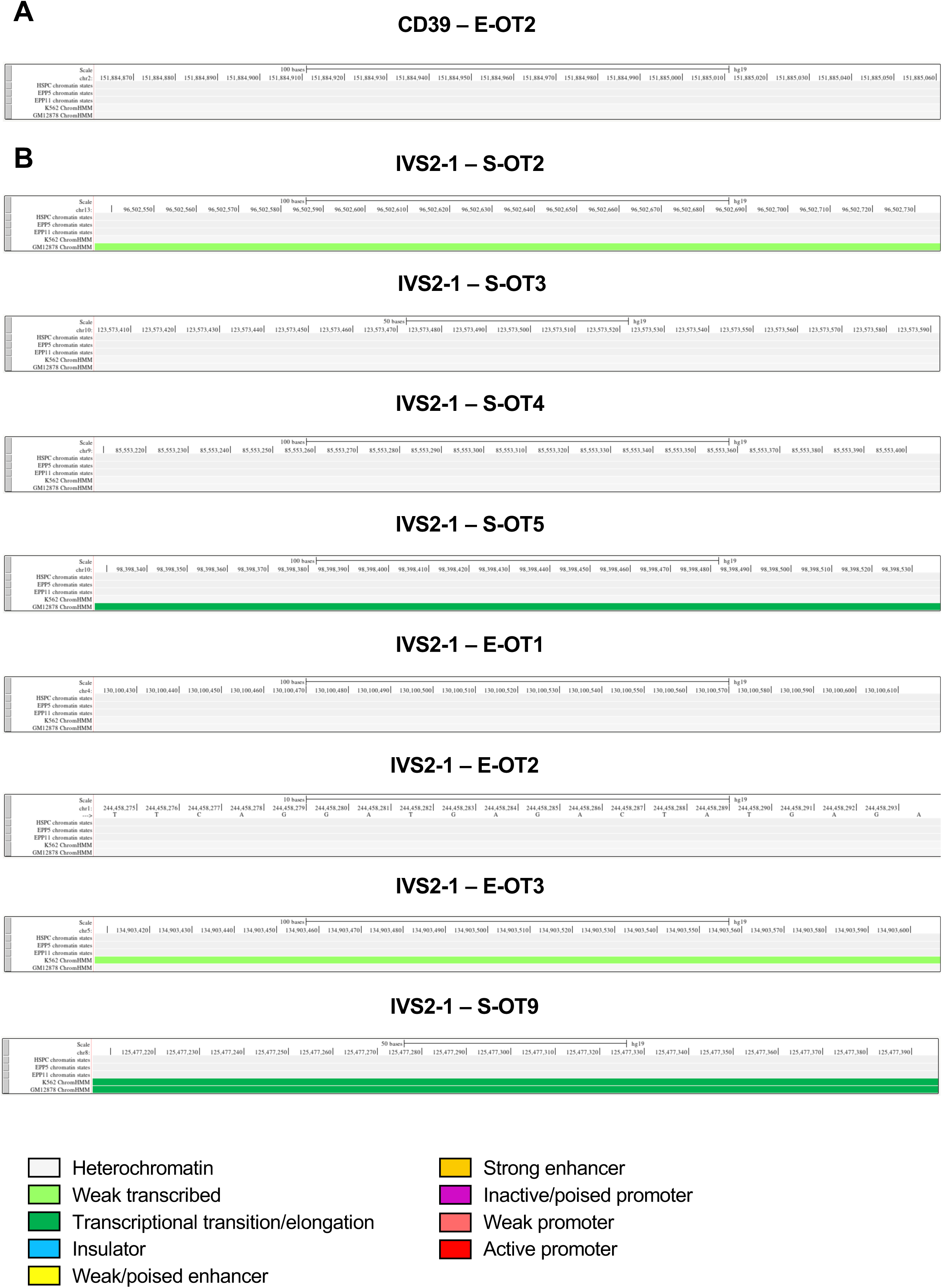
Chromatin state of off-target loci. Analysis of the chromatin states of CD39 (**A**) and IVS2-1 (**B**) validated off-target sites in primary human HSPCs, HSPC-derived early (EPP5) and late (EPP11) erythroid precursors, and in erythroid (K562) and granulo-monocytic (GM12878) cell lines (UCSC datasets).

**Supplementary Figure 7.**
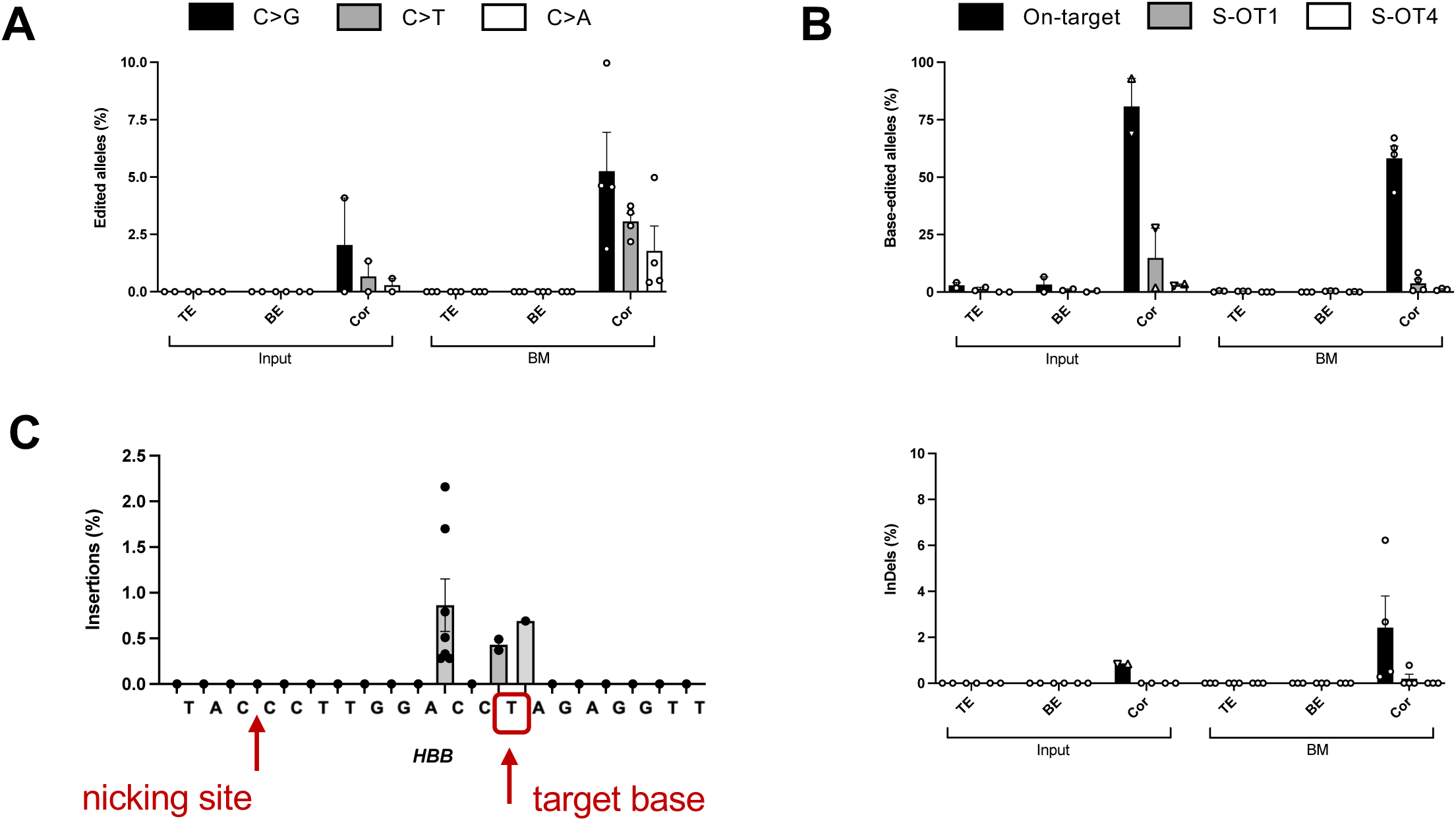
On- and off-target editing *in vivo*. **A.** Frequency of cytosine bystander editing (b3, identified in **Supplemental Figure 2**) as calculated after deep sequencing of input and BM human CD45^+^ cells derived from mice transplanted with control and edited BT-HSPCs. Data are expressed as mean±SEM (Input: n=2, BT-TE/BE: n=3, BT-cor: n=4). **B.** Frequency of base-edited alleles (upper panel) and InDels (lower panel) as calculated after deep sequencing of on-target site, S-OT1 and S-OT4 in input populations and BM human CD45^+^ cells derived from mice transplanted with control and edited BT-HSPCs. Data are expressed as mean±SEM (Input: n=2, BT-TE/BE: n=3, BT-cor: n=4). The editing frequency in the input was calculated in cells cultured in the HSPC medium (▼) or in liquid erythroid cultures (▲). **C.** Frequency and distribution of insertions across the CD39 gRNA1 protospacer. Each dot corresponds to the frequency of an insertion event identified in BM human CD45^+^ cells derived from mice transplanted with edited BT-HSPCs. Data are expressed as mean±SEM.

---

**Supplemental Table 1.**
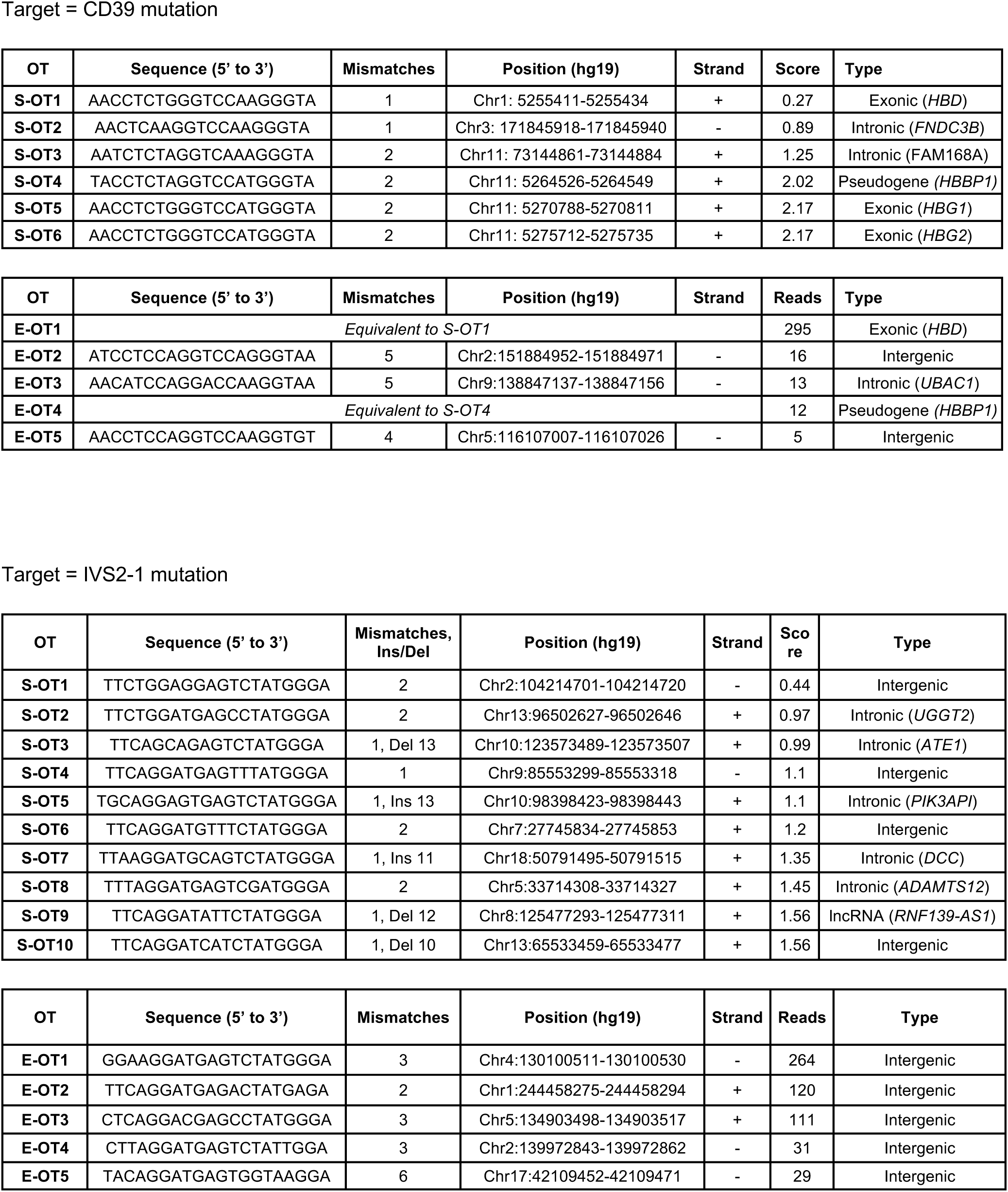
*In silico* and empirically nominated off target sites.

**Supplemental Table 2.**
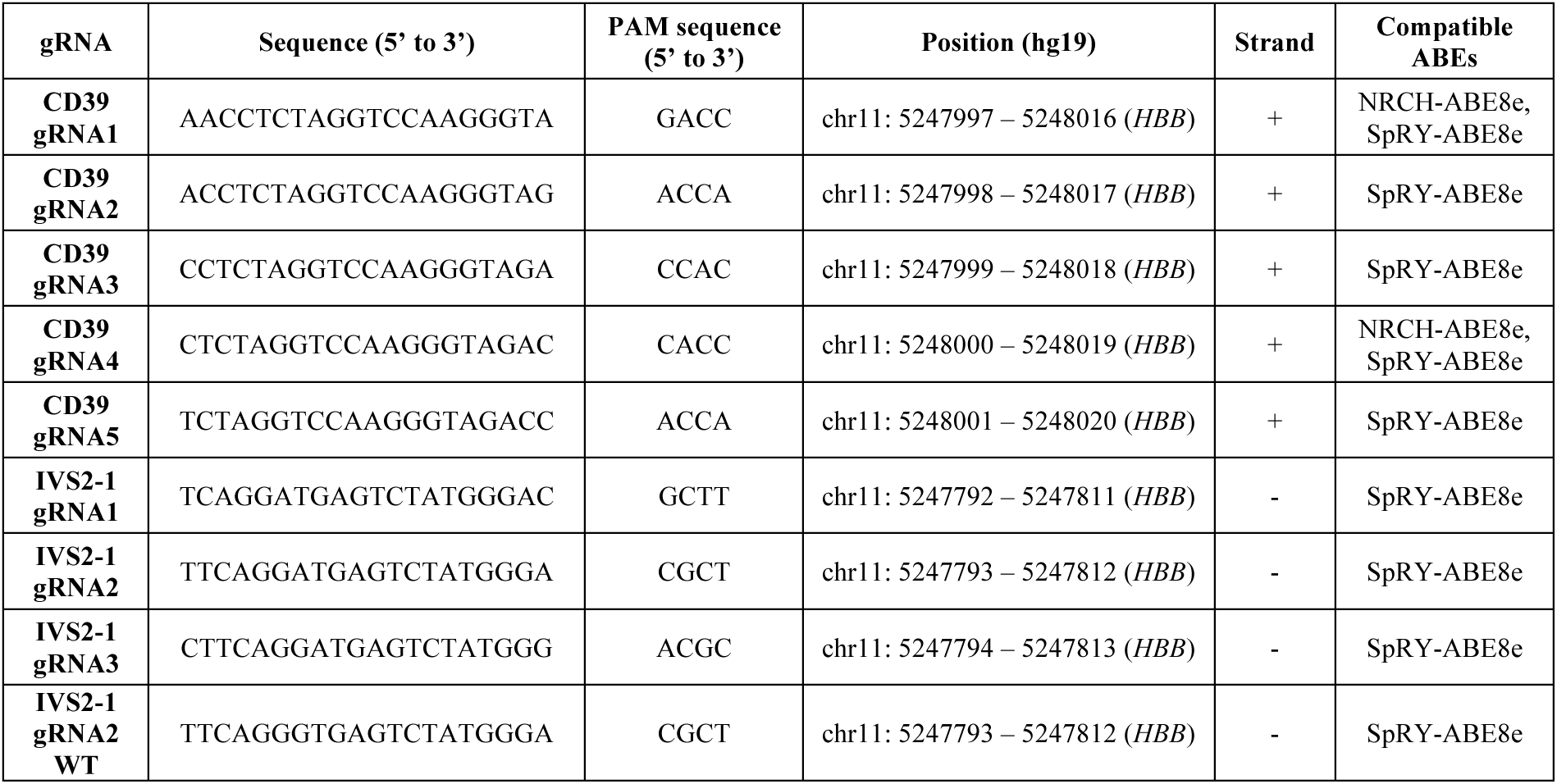
gRNA target sequences.

**Supplemental Table 3.**
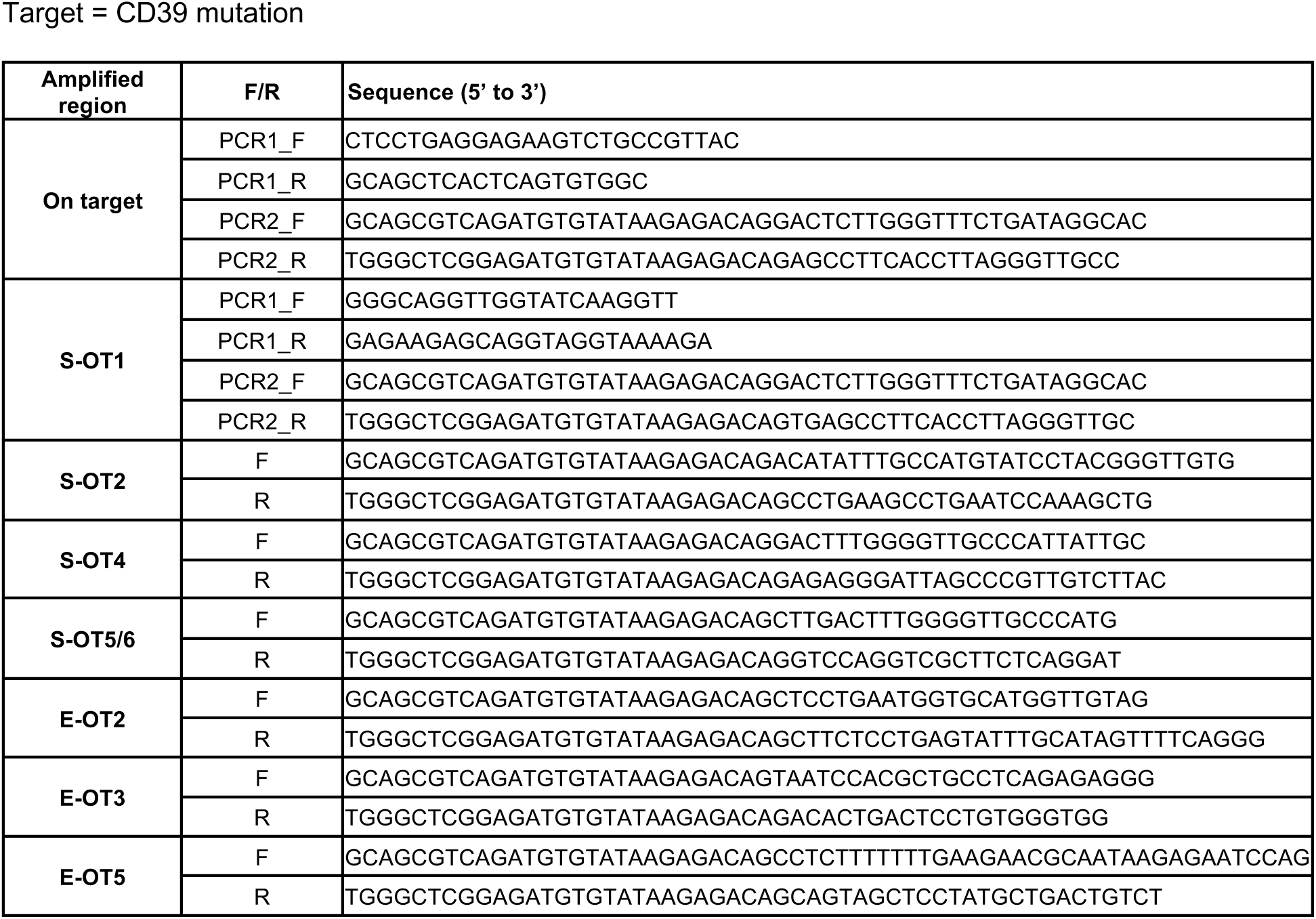
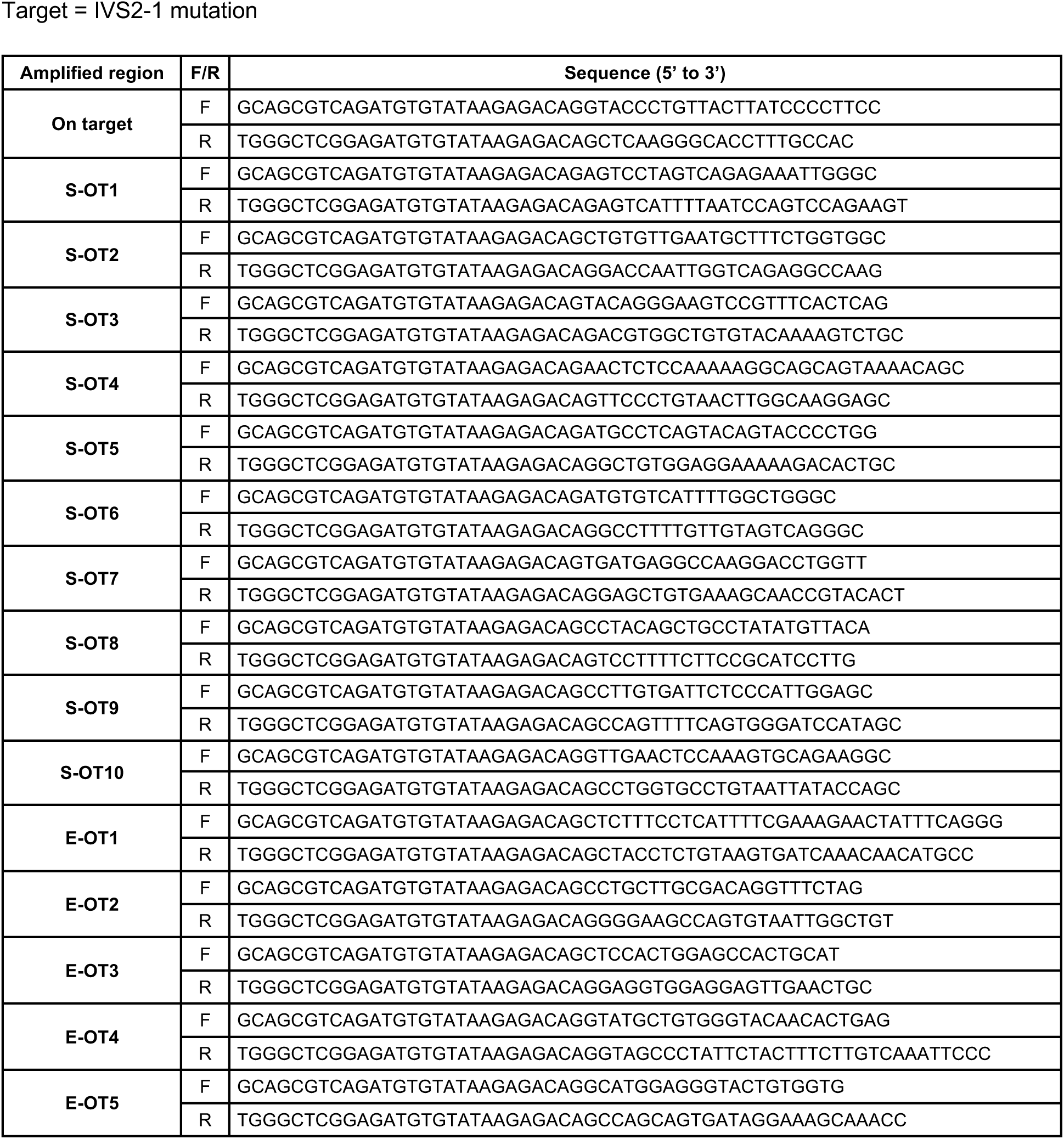
PCR primers to amplify on-target and off-target sites.

**Supplemental Table 4.**
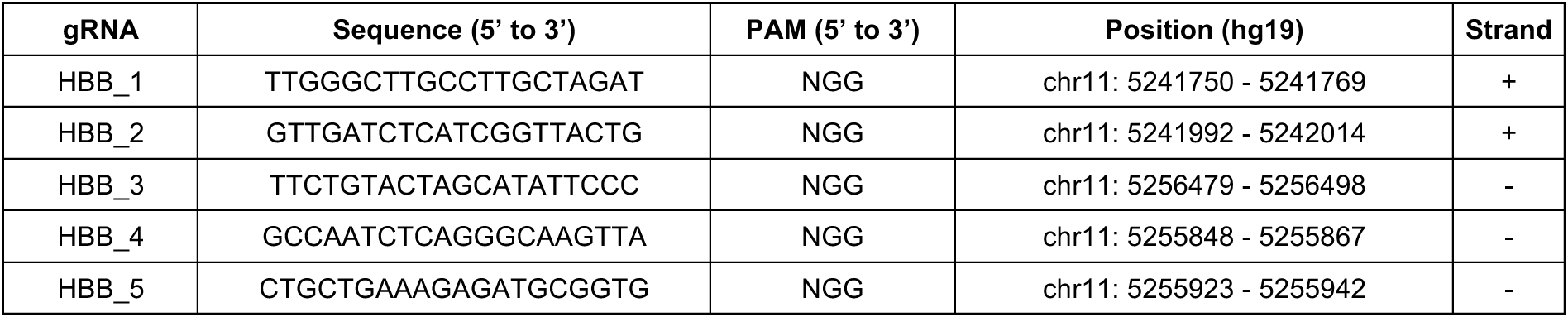
gRNAs used for Cas9-enrichment library preparation.

The following investigators contributed to NaThalY, composing the **Consortium NaThalY** (listed in alphabetic order): Kokou-Placide Agbo-Kpati, CH Marne la vallée, Paris; Nathalie Aladjidi, Hôpital Pellegrin, Bordeaux; Oula Alhindy, Hôpital Henri Mondor, Aurillac; Slimane Allali, Hôpital Necker, Paris; Shanti Ame, Hôpital Civil, Strasbourg; Jean-Benoit Arlet, Hôpital Européen George Pompidou, Paris; Corinne Armari, CHU Grenoble, Grenoble; Sai Atmani, Hôpital Laennec, Creil; Dora Bachir, Hôpital Henri Mondor, Créteil; Catherine Badens, Hôpital de la Timone, Marseille ; Melissa Barbati, Hôpital Jeanne de Flandre, Lille; Sandrine Baron-Joly, CHU de Nîmes, Nimes; Fiorenza Barraco, Hôpital Edouard Herriot, Lyon; Pablo Bartolucci, Hôpital Henri Mondor, Créteil; Sophie Bayart, CHU Rennes, Rennes; Marie Belloy, CHIC Robert Ballanger, Aulnay-Sous-Bois; Malika Benkerrou, Hôpital Robert Debré, Paris; Julie Berbis, Hôpital de La Timone, Marseille; Ana Berceanu, CHU Besancon, Besançon; Claire Berger, CHU Nord Saint Etienne, Saint-Etienne; Emmanuelle Bernit, CHU de Pointe à Pitre, Guadeloupe; Ivan Bertchansky, CHU Lapeyronie, Montpellier; Yves Bertrand, IHOP, Lyon; Anne Besancon-Bergelin, CH Le Mans, Le Mans; Gilles Blondin, CHIC Quimper, Quimper; Nicolas Boissel, Hôpital Saint Louis, Paris; Jacinthe Bonneau-Lagacherie, CHU Rennes, Rennes; Emmanuelle Bourgeois-Petit, Hôpital Saint Vincent de Paul, Lille; Jean-François Brasme, CHU d’Angers, Angers; Valentine Brousse, Hôpital Robert Debré, Paris; Giovanna Cannas, Hôpital Edouard Herriot, Lyon; Benjamin Carpentier, Hôpital Saint Vincent de Paul, Lille; Marie-Pierre Castex, CHU de Toulouse, Toulouse; Marina Cavazzana, Hôpital Necker, Paris; Abdourahim Chamouine, CH Mayotte, Mayotte; Christelle Chantalat-Auger, Hôpital Bicêtre, Le Kremlin Bicêtre; Thibault Comont, IUCT, Toulouse; Mohamed Conde, Hôpital de La Source, Orléans; Marie-Laure Couec, Hôpital enfant-adolescent, Nantes; Pierre Cougoul, IUCT, Toulouse; Jean-Hugues Dalle, Hôpital Robert Debré, Paris; Audrey David, CHU Nord Saint Etienne, Saint-Etienne; Sébastien De Almeida, CHU de Toulouse, Toulouse; Gonzalo De Luna, Hôpital Henri Mondor, Créteil; Marianne De Montalembert, Hôpital Necker, Paris; Benjamin De Sainte Marie, Hôpital de la Timone, Marseille; Aurélie Desbree, CHU Nord Saint Etienne, Saint-Etienne; Marie Detrait, Hôpital Brabois, Vandoeuvre Les Nancy; Alain Devidas, Centre Hospitalier Sud Francilien, Corbeil-Essonnes; Nathalie Dhedin, Hôpital Saint Louis, Paris; Judith Diakiesse, CH François Quesnay, Mantes La Jolie; Pierre-Yves Dides, CH de GRASSE, Grasse; Jean Donadieu, Hôpital Armand Trousseau, Paris; Antoine Dossier, Hôpital Bichat, Paris; Grégoire Ducoux, Hôpital Edouard Herriot, Lyon; Pierre Duffau, Groupe Hospitalier Saint André, Bordeaux; Fabienne Dulieu, CHU Lenval, Nice; Cecile Dumesnil, Hôpital Charles Nicolle, Rouen; Julie Durin, CHIC Robert Ballanger, Aulnay-Sous-Bois; Sylvie Fasola, Hôpital Armand Trousseau, Paris; Benoit Faucher, Hôpital de la Timone, Marseille; Bruno Filhon, Hôpital Charles Nicolle, Rouen; Elena Fois, Hôpital Henri Mondor, Créteil; Edouard Forcade, Hôpital Pellegrin, Bordeaux; Frederic Galacteros, Hôpital Henri Mondor, Créteil; Loïc Garcon, CHU Sud Amiens, Amiens; Reda Garidi, CHU Sud Amiens, Amiens; Nathalie Garnier, IHOP, Lyon; Jean-Baptiste Gaultier, CHU Nord Saint Etienne, Saint-Etienne; Alexandra Gauthier, IHOP, Lyon; Justine Gellen-Dautremer, Hôpital La Miletrie, Poitiers; Fanny Gonzales, Hôpital Jeanne de Flandre, Lille; François Gouraud, Grand Hôpital de l’Est Francilien, Meaux; Stephanie Gourdon, CHU La Réunion, La Réunion; Isabelle Guichard, CHU Nord Saint Etienne, Saint-Etienne; Henri Guillet, Hôpital Henri Mondor, Créteil; Corinne Guitton, Hôpital Bicêtre, Le Kremlin Bicêtre; Anoosha Habibi, Hôpital Henri Mondor, Créteil; Nabil Heouaine, CH de Dunkerque, Dunkerque; Laurent Holvoet, Hôpital Robert Debré, Paris; Arnaud Hot, Hôpital Edouard Herriot, Lyon; Yoann Huguenin, Hôpital Pellegrin, Bordeaux; Ghislaine Ithier, Hôpital Robert Debré, Paris; Estelle Jean, Hôpital de la Timone, Marseille; Laure Joseph, Hôpital Necker, Paris; Annie Kamdem, CHI Créteil, Créteil; Jean-Michel Karsenti, CHU l’Archet, Nice; Kamila Kebaili, IHOP, Lyon; Bérengère Koehl, Hôpital Robert Debré, Paris; Florence Lachenal, Hôpital Pierre Oudot, Bourgoin-Jallieu; Mathieu Lacou, CHU de Nantes, Nantes; Agnes Lahary, Hôpital Charles Nicolle, Rouen; Muriel Lalande, CHU Lapeyronie, Montpellier; Anne Lambilliotte, Hôpital Jeanne de Flandre, Lille; Thierry Leblanc, Hôpital Robert Debré, Paris; Agnes Lehnert, CHAM, Montargis; Guy Leverger, Hôpital Armand Trousseau, Paris; Valérie Li Thiao Te, CHU Nord Amiens, Amiens; François Lionnet, Hôpital Tenon, Paris; Bruno Lioure, Hôpital de Hautepierre, Strasbourg; Marie-Josée Lucchini, CH de la Miséricorde, Ajaccio; Laurence Lutz, Hôpital de Hautepierre, Strasbourg; Abdallah Maakaroun, CH Jacques-Coeur, Bourges; Alexis Magda, Hôpital de La Source, Orléans; Perrine Mahe, CHU Lapeyronie, Montpellier; Julie Maitre, Hôpital de La Source, Orléans; Caroline Makowski, CHU Grenoble, Grenoble; Aurore Malric, Hôpital Delafontaine, Saint-Denis; Corina Manoli, Hôpital Européen George Pompidou, Paris; Agathe Masseau, Hôpital Hôtel-Dieu, Nantes; Caroline Masserot-Lureau, Grand Hôpital de l’Est Francilien, Meaux; Catherine Mathey, CH Aix en Provence, Aix En Provence; Suzanne Mathieu, Hôpital Brabois, Vandoeuvre Les Nancy; Sarah Mattioni, Hôpital Tenon, Paris; Benoit Meunier, Hôpital Necker, Paris; Gerard Michel, Hôpital de la Timone, Marseille; Florence Missud, Hôpital Robert Debré, Paris; Despina Moshous, Hôpital Necker, Paris; Robert Navarro, CHU Lapeyronie, Montpellier; Benedicte Neven, Hôpital Necker, Paris; Stéphanie Ngo, Hôpital Charles Nicolle, Rouen; Stanislas Nimubona, Hôpital Pontchaillou, Rennes; Laure Nizery, Hôpital Delafontaine, Saint-Denis; Placide Nyombe, CHU La Réunion, La Réunion; Moïse Ondja, CH Vitry le François, Vitry-Le-François; Corentin Orvain, CHU d’Angers, Angers; Eric Oziol, CH de Béziers, Béziers; Marlène Pasquet, CHU de Toulouse, Toulouse; Régis Peffault De La Tour, Hôpital Saint Louis, Paris; Charline Pegon, CHU Estaing, Clermont-Ferrand; Sophie Pertuisel, CHU Rennes, Rennes; Charlotte Petitdidier, Centre Hospitalier Sud Francilien, Corbeil-Essonnes; Aurélie Phulpin, Hôpital Brabois, Vandoeuvre Les Nancy; Christophe Piguet, Hôpital Dupuytren, Limoges; Geneviève Plat-Wilson, CHU de Toulouse, Toulouse; Claire Pluchart, American Memorial Hospital, Reims; Cécile Pochon, Hôpital Brabois, Vandoeuvre Les Nancy; Maryline Poiree, CHU l’Archet, Nice; Corinne Pondarre, CHI Créteil, Créteil; Nathalie Ravet, CH Marne la Vallée, Paris; Yves Reguerre, CHU La Réunion, La Réunion; Cécile Renard, IHOP, Lyon; Claire Reynes, CH Annecy Genevois, Annecy; Fanny Rialland, CHU de Nantes, Nantes; Jean-Antoine Ribeil, Hôpital Necker, Paris; Marie Robin, Hôpital Saint Louis, Paris; Pierre-Simon Rohrlich, CHU l’Archet, Nice; Aphaia Roussel, Hôpital de La Timone, Marseille; Lara Sabbagh, Hôpital Henri Mondor, Aurillac; Alexandra Salmon, Hôpital de Hautepierre, Strasbourg; Paul Saultier, Hôpital de La Timone, Marseille; Jean-Marc Schneider, CH Bel-Air, Metz-Thionville; Maud Schoeny, CH Saint Dizier, Saint Dizier; JILL Serre, Hôpital Clocheville, Tours; Pauline Simon, Hôpital Saint Jacques, Besançon; Anne Sirvent, CHU Lapeyronie, Montpellier; Borhane Slama, CH Henri Duffaut, Avignon; Gérard Socie, Hôpital Saint Louis, Paris; Bertrand Soto, Hôpital Simone Veil, Troyes; Alexandra Spiegel, Hôpital de Hautepierre, Strasbourg; Katia Stankovic, Hôpital Tenon, Paris; Jean-Louis Stephan, CHU Nord Saint Etienne, Saint-Etienne; Arthur Sterin, Hôpital de la Timone, Marseille; Dominique Steschenko, Hôpital Brabois, Vandoeuvre Les Nancy; Cecile Stoven, CHU La Réunion, La Réunion; Laure Swiader, Hôpital de la Timone, Marseille; Sarah Szepetowski, Hôpital de La Timone, Marseille; Faustine Tardy, CHU Nord Saint Etienne, Saint-Etienne; Mélissa Taylor, Hôpital Necker, Paris; Caroline Thomas, CHU de Nantes, Nantes; Isabelle Thuret, Hôpital de la Timone, Marseille; Mohamed Touati, Hôpital Dupuytren, Limoges; Capucine Trochu, CH Roubaix, Roubaix; Jean-Baptiste Valentin, Hôpital Trousseau, Tours; Amélie Vareliette, Hôpital de Brest, Brest; Bruno Varet, Hôpital Necker, Paris; Emilie Virot, Hôpital Edouard Herriot, Lyon; Agathe Wautier-Rascalou, CHU de Nîmes, Nîmes; Alienor Xhaard, Hôpital Saint Louis, Paris; Karima Yakouben, Hôpital Robert Debré, Paris; Marion Yvert, Hôpital Clocheville, Tours; Ferielle Zenchri, Hôpital Bicêtre, Le Kremlin Bicêtre.

